# PSTPIP2 ameliorates aristolochic acid nephropathy by suppressing interleukin-19-mediated neutrophil extracellular trap formation

**DOI:** 10.1101/2023.07.24.550330

**Authors:** Changlin Du, Jiahui Dong, Chuanting Xu, Pengcheng Jia, Na Cai, Qi Wang, Zhenming Zhang, Lingfeng Jiang, Wei Jiang, Rui Feng, Jun Li, Cheng Huang, Taotao Ma

## Abstract

Aristolochic acid nephropathy (AAN) is a progressive kidney disease caused by herbal medicines. Previously, we found that proline–serine–threonine phosphatase interacting protein 2 (PSTPIP2) and neutrophil extracellular traps (NETs) play important roles in kidney injury and immune defense, respectively; however, the mechanism of AAN regulation by PSTPIP2 and NETs remains unclear. We found that renal tubular epithelial cell (RTEC) apoptosis, neutrophil infiltration, and inflammatory factor and NET production were increased in a mouse model of AAN, while PSTPIP2 expression was low. Conditional knock-in of PSTPIP2 in mouse kidneys inhibited cell apoptosis, reduced neutrophil infiltration, suppressed the production of inflammatory factors and NETs, and ameliorated renal dysfunction. In contrast, restoring normal PSTPIP2 expression promoted kidney injury. *In vivo,* the use of Ly6G-neutralizing antibody to remove neutrophils and peptidyl arginine deiminase 4 (PAD4) inhibitors to prevent NET formation reduced apoptosis, thereby alleviating kidney injury. *In vitro*, damaged RTECs released interleukin-19 (IL-19) via the PSTPIP2/nuclear factor (NF)-κB pathway and induced NET formation via the IL-20Rβ receptor. Concurrently, NETs promoted the apoptosis of damaged RTECs. PSTPIP2 affected NET formation by regulating IL-19 expression via inhibition of NF-κB pathway activation in RTECs, inhibiting their apoptosis and reducing kidney damage.

## Introduction

The World Health Organization has estimated that 80% of the global population use traditional medicine (1). However, the pharmacologically active components of some traditional Chinese herbal medicines are associated with adverse effects (2). Aristolochic acids (AAs) are found in herbs of the genus *Aristolochia* and are well known for their nephrotoxicity and carcinogenicity (3). Notably, 8-Methoxy-6-nitrophenanthro-(3,4-d)-1,3-di-oxolo-5-carboxylic acid (AAI) is a major AA (4). Aristolochic acid nephropathy (AAN) refers to the deterioration in kidney function associated with toxic AAs (5). AAN is one of the most serious complications associated with the use of traditional Chinese medicines, and cases of the disease have been reported globally (6). Both mice and rats are susceptible to AA toxicity, allowing *in vivo* studies on the mechanisms of AA-induced renal toxicity. Acute renal failure with tubular necrosis can be induced in these model animals by a single high dose of AA (7). Currently, there is no effective treatment for AAN; thus, further investigation is required to elucidate the mechanisms of AAI-induced acute kidney injury.

Under conditions of persistent injury, signals from the damaged cells activate inflammatory cells, which release proinflammatory cytokines and stimulate the apoptosis of the injured cells. Cell communication is an essential component of mammalian development and homeostasis, ensuring fast and efficient responses to alterations or threats within the host cell (8). Inflammation is the body’s protective response against external threats and promotes the programmed death of damaged cells to maintain homeostasis, promote the adaptation of tissue to harmful conditions, and restore tissue function (9). Several studies have shown that inflammatory cell infiltrates, excessive injury, and death of renal tubular epithelial cells (RTECs) characterize the acute phase of AAN (10–12). Neutrophils, a major inflammatory cell type, infiltrate the kidney in the early stages of AAN, leading to injury. However, it remains unclear how RTEC injury promotes neutrophil activation and how this stimulates the apoptosis of damaged RTECs in AAN.

In response to certain stimuli, neutrophils produce neutrophil extracellular traps (NETs)—web-like structures composed of decondensed DNA, histones, and neutrophil granule proteins—through which neutrophils cause tissue damage and interact with other cells. NETs can exacerbate tissue damage via several mechanisms, including promotion of vascular occlusion(13), sterile inflammation(14), and immune balance disruption. Saisorn *et al.* (15) reported that increased lupus activity may exacerbate acute kidney injury (AKI) through NETs and NET-induced apoptosis. NETs are important for immune defense; however, their role in mediating apoptosis remains controversial. Lv *et al.* (16) reported that NET formation is a major cytotoxic factor causing lung epithelial apoptosis and cytoskeletal destruction. Shen *et al.* (17) also found that NETs promoted trophoblast apoptosis in gestational diabetes mellitus. However, the role of NETs and their potential regulatory mechanisms in AAN remain unclear.

In this study, we aimed to determine whether NETs play a role in the survival of injured RTECs and elucidate the underlying mechanism. We provide evidence that RTECs release interleukin (IL)-19 via a PSTPIP2/nuclear factor (NF)-κB-dependent process after AA stimulation, and initiate a feed-forward mechanism involving IL-20Rβ-dependent NET formation. Furthermore, NETs were found to induce apoptosis in AAI-treated RTECs. Overall, tubular epithelial cell-to-neutrophil-to-tubular epithelial cell circulation via NET transfer promoted renal inflammation and apoptosis in AAN. Our elucidation of the interaction between RTECs and neutrophils has potential clinical implications in the treatment of AAN.

## Results

### *PSTPIP2* expression decreased in the AAI-induced acute AAN model *in vivo* and AAI-treated RTECs *in vitro*

To investigate the natural course of the renal response to AAI, kidney tissues were harvested on days 1, 3, and 5 after AAI administration (Figure S1A). Renal impairment, based on elevated serum creatinine (Cr) and blood urea nitrogen (BUN) levels, significantly increased on days 3 and 5 (Figure S1B and C). Consistent with the severity of tubular damage observed in hematoxylin and eosin (H&E)-stained sections (Figure 1A), levels of the biomarker of tubular damage, KIM-1, markedly increased on days 3 and 5 as well, as detected by western blotting and immunocytochemistry (Figure S1D and E). The expression of PSTPIP2 in the kidney tissue of the AAI-treated group was significantly lower than that in the vehicle group (Figure 1B and C). Double immunofluorescence (IF) staining showed that PSTPIP2 and E-cadherin were co-localized (Figure 1D). E-cadherin is an important epithelial cell marker, suggesting that PSTPIP2 is expressed in RTECs. Localization analysis demonstrated that AAI treatment decreased the expression of PSTPIP2 in RTECs. RTECs were the main site of cell injury during AAI nephrotoxicity. We used AAI to induce AKI in mRTECs *in vitro*. We first evaluated appropriate treatment conditions and found that a 20-h treatment with AAI did not induce apoptosis in RTECs but increased cell injury; thus, this condition was used for our experiments (Figure S1F–H). Western blot analysis (Figure 1E and F) and IF staining (Figure 1G and H) showed that the expression of KIM-1 was significantly upregulated and that of PSTPIP2 was downregulated in AAI-treated mRTECs compared to that in the vehicle-treated group. Collectively, these results indicate that PSTPIP2 expression decreased in both mice and mRTECs when treated with AAI.

**Figure 1.**
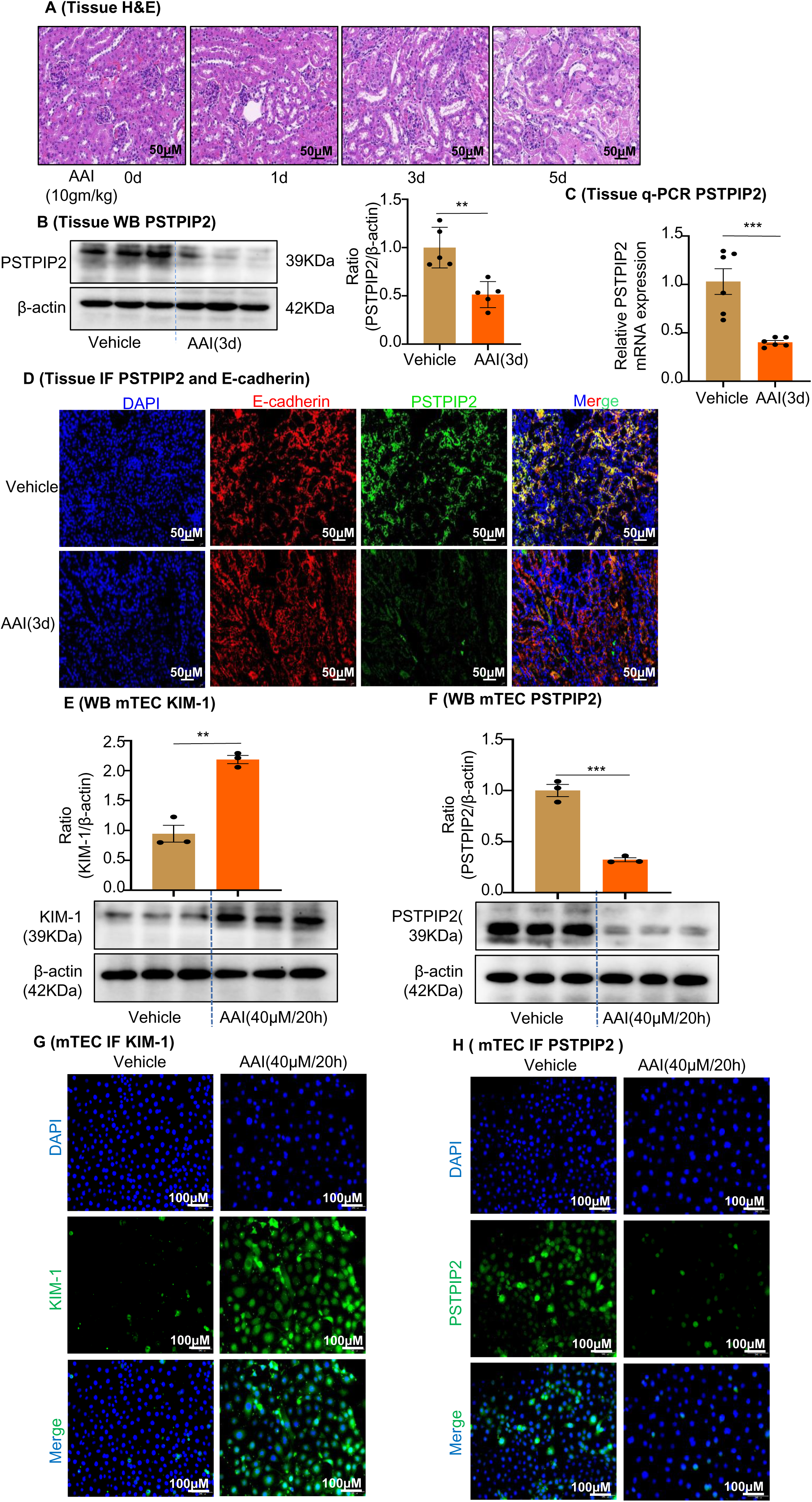
*PSTPIP2* expression was decreased in the aristolochic acid I (AAI) induced acute aristolochic acid nephropathy model *in vivo* and injured mouse renal tubular epithelial cells (mRTECs) *in vitro*. **(A)** Representative H&E-stained images of kidney sections. Kidneys were isolated on day 0, 1, 3, and 5 after AAI treatment (n = 6 per group). Scale bar, 50 μm. **(B)** Western blot analysis of PSTPIP2 expression in kidney tissue lysates, which were isolated on day 3 after AAI treatment. β-actin was used as a loading control. Quantification of the PSTPIP2/β-actin ratio (n = 5 per group). **(C)** Real-time PCR analysis of *PSTPIP2* expression in kidney tissue, which were isolated on day 3 after AAI treatment. β-actin was used as a loading control (n = 6 per group). **(D)** Immunofluorescence (IF) staining of PSTPIP2 (green) and E-cadherin (red, marker for epithelial cells) in kidney tissues. DAPI was used for nuclear staining (n = 6 per group). Scale bar, 50 μm. **(E)** Protein levels of kidney injury molecule-1 (KIM-1) in mRTECs treated with AAI (40 μM/20 h) based on western blotting. Quantification of the KIM-1/β-actin ratio (n = 3). **(F)** Protein levels of PSTPIP2 in mRTECs treated with AAI (40 μM/20 h) analyzed by western blotting. Quantification of the PSTPIP2/β-actin ratio (n = 3). **(G)** Protein levels of KIM-1 in mRTECs treated with AAI (40 μM/20 h) analyzed by IF (n = 3). Scale bar, 100 μm. **(H)** Protein levels of PSTPIP2 in mRTECs treated with AAI (40 μM/20 h) analyzed by IF (n = 3). Scale bar, 100 μm. Data are presented as the mean ± SEM of three to six biological replicates per condition. Each dot represents a sample. Statistically significant differences were determined by an independent sample *t*-test (B,C,E,F). *P < 0.05, **P < 0.01, ***P < 0.001, ns: non-significant. **Figure 1—source data 1** **Figure 1—source data 2** ***PSTPIP2* expression was decreased in the aristolochic acid I (AAI)-induced acute aristolochic acid nephropathy model *in vivo* and injured mouse renal tubular epithelial cells (mRTECs) *in vitro*.**

### Changes in renal *PSTPIP2* expression could affect AAI-induced kidney injury and apoptosis

To determine the role of renal PSTPIP2 levels in AAI-induced AAN, we established a PSTPIP2 conditional knock-in (*PSTPIP2*-cKI) mouse model by CRISPR/Cas mediated genome engineering (Figure 2A); PSTPIP2 was indeed overexpressed in the RTECs. PSTPIP2^Flox/Flox^ (FF) mice served as controls. All mice were genotyped using PCR (Figure 2B). IF and PCR analyses showed that PSTPIP2 was successfully knocked in in the mouse kidneys (Figure 2C and D). Renal PSTPIP2 overexpression drastically ameliorated renal damage when treated with AAI, based on a histological analysis with H&E staining (Figure 2E). Similarly, Cr and BUN levels were remarkably attenuated in AAI-treated *PSTPIP2*-cKI mice compared to those in AAI treated FF mice (Figure 2F and G). In addition, western blot and PCR analysis showed that KIM-1 expression was significantly upregulated in AAI-treated FF mice, while that in AAI-treated *PSTPIP2*-cKI mice was reduced (Figure 2H and I).

**Figure 2.**
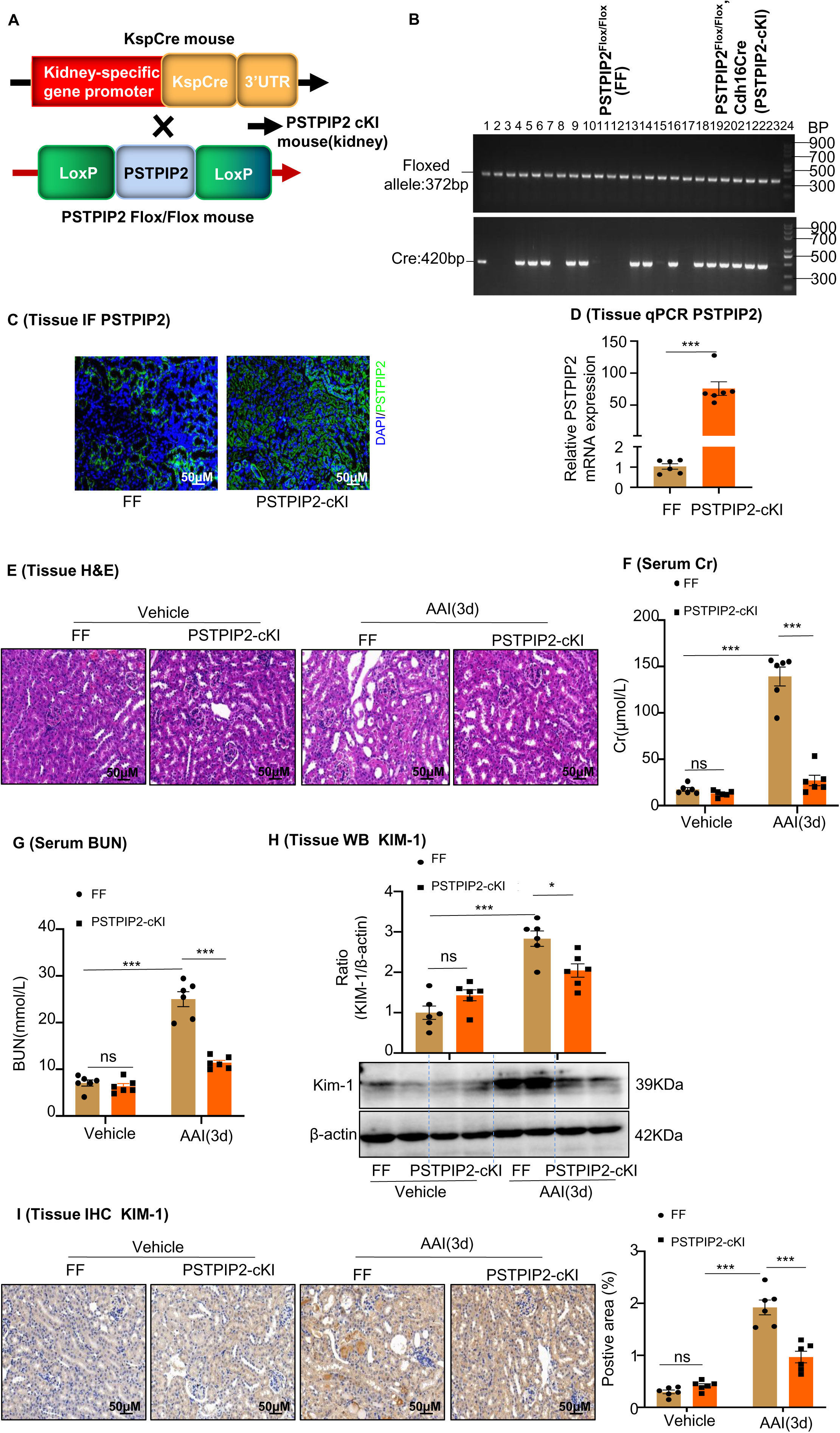
*PSTPIP2*-cKI attenuates aristolochic acid I (AAI)-induced kidney damage. **(A)** Scheme illustrating the genetic approach for generating *PSTPIP2* conditional knock-in (cKI) mice. **(B)** Genotypes of *PSTPIP2*-cKI mice were confirmed using PCR. The floxed allele showed a 372 bp sized band, and the Cre allele was 420 bp; 15 *PSTPIP2*^Flox/Flox^-Cdh16Cre (or *PSTPIP2*-cKI) mice (#1, #4-6, #8, #9, #13, #14, #16, #18-23) and 9 PSTPIP2^Flox/Flox^ (or FF) mice (#2, #3, #7, #10-12, #15, #17, #24) are shown (n = 24). **(C)** Immunofluorescence (IF) staining of *PSTPIP2* in FF (n = 6) and *PSTPIP2*-cKI (n = 6) mice. Scale bar, 50 μm. **(D)** mRNA level of *PSTPIP2* in FF (n = 6) and *PSTPIP2*-cKI (n = 6) mice assessed by real-time PCR. **(E)** H&E staining of kidneys from FF (n = 6) and *PSTPIP2*-cKI (n = 6) mice treated with AAI. Scale bar, 50 μm. **(F and G)** Serum creatinine (Cr) and blood urea nitrogen (BUN) assay. Serum Cr and BUN showed that *PSTPIP2* overexpression prevented renal dysfunction in AAI-induced nephropathy (n = 6 per group). **(H)** Expression of kidney injury molecule-1 (KIM-1) protein in kidney tissues from FF and *PSTPIP*2-cKI mice treated with vehicle and AAI examined by western blotting (n = 6 per group). **(I)** Representative immunohistochemical (IHC) staining and KIM-1 levels in kidney tissues (n = 6 per group). Scale bar, 50 μm. Data are presented as the mean ± SEM of six biological replicates per condition. Each dot represents a sample. Statistically significant differences were determined by an independent sample *t*-test (D) and one-way ANOVA followed by Tukey’s post hoc test (F–I). *P < 0.05, **P < 0.01, ***P < 0.001, ns: non-significant. **Figure 2—source data 1** **Figure 2—source data 2** ***PSTPIP2*-cKI attenuates aristolochic acid I (AAI)-induced kidney damage.**

RTEC apoptosis is a common histopathological feature of AAN. TUNEL staining revealed that TUNEL^+^ cells increased in AAI-treated mice compared to the control mice (Figure S2A). As caspase-3 plays a central role in the execution phase of apoptosis, we detected its activity and the expression of its activated form, cleaved caspase-3, in mouse kidney tissues. The activity of caspase-3 and level of cleaved caspase-3 were markedly increased in mice treated with AAI (Figure S2B–D). Conditional knock in of PSTPIP2 significantly decreased TUNEL^+^ cell numbers compared with those in the AAI-treated WT group (Figure S2E). Furthermore, the activity of caspase-3 and level of cleaved caspase-3 were significantly downregulated in AAI-treated *PSTPIP2*-cKI mice compared with those in AAI-treated FF mice (Figure S2F–H). These results indicate that PSTPIP2 has a protective effect against AAI-induced AAN.

To further examine the functional importance of PSTPIP2, we silenced PSTPIP2 expression using an AAV9-packaged *PSTPIP2* shRNA plasmid in *PSTPIP2*-cKI mice (Figure 3A, C–E). Fluorescence-labelled AAV9–*PSTPIP2* was localized to the kidneys (Figure 3B). We found that a reduction in *PSTPIP2* expression increased serum BUN and Cr levels, along with renal damage (Figure 3F–H). Furthermore, immunohistochemical (IHC) staining and western blotting showed that the reduction of *PSTPIP2* expression led to the elevated expression of KIM-1 in AAI-induced *PSTPIP2*-cKI mice (Figure 3J and I). In addition, the reduction of *PSTPIP2* expression significantly increased TUNEL^+^ cell numbers, caspase-3 activity, and cleaved caspase-3 levels compared to those in AAI-treated *PSTPIP2*-cKI mice (Figure 3K–N). These findings indicate that PSTPIP2 plays a key role in AAI-induced renal injury and apoptosis.

**Figure 3.**
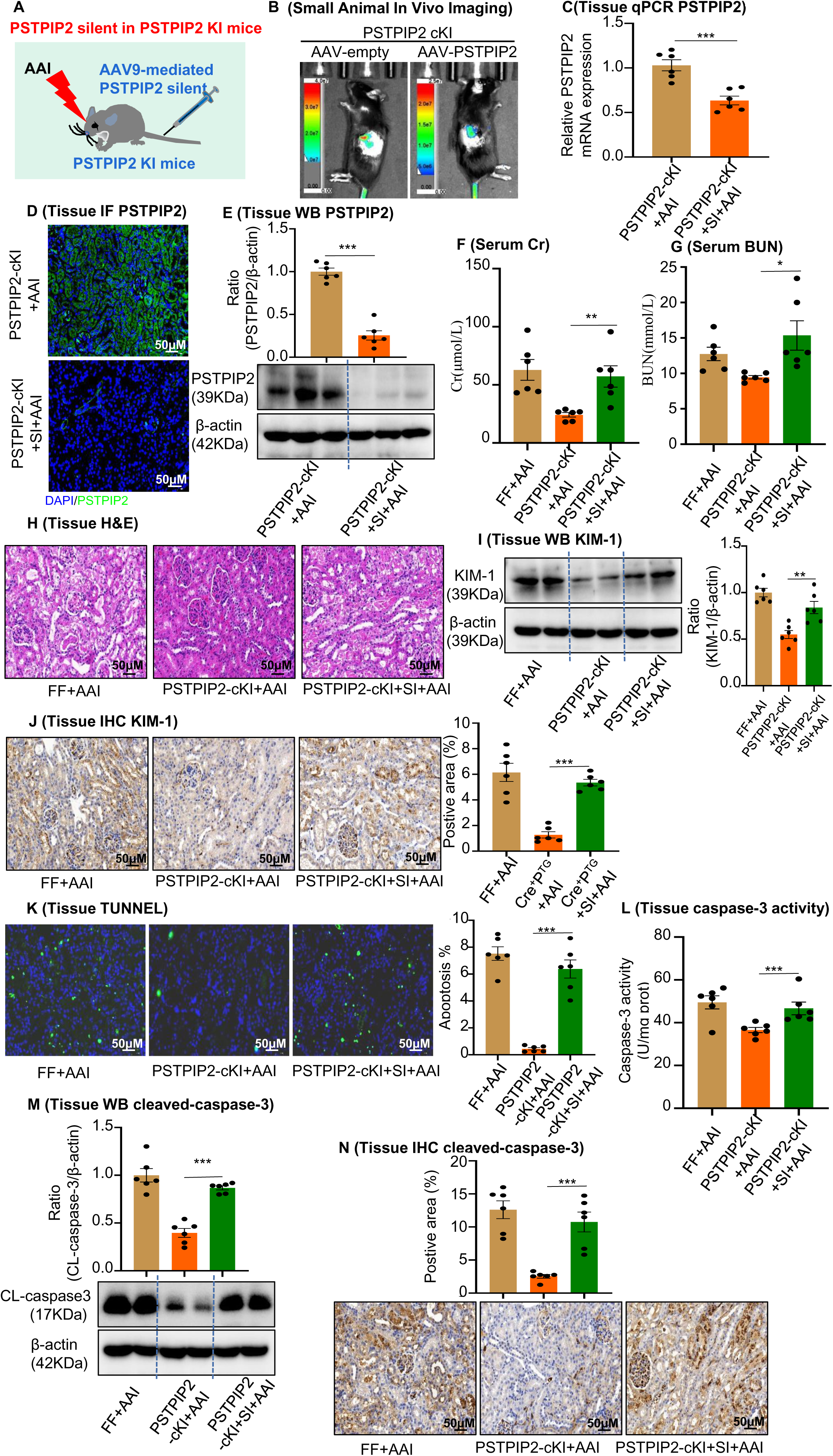
AAV9-mediated *PSTPIP2* restoration re-induces renal injury and kidney apoptosis in *PSTPIP2*-cKI mice. **(A)** Scheme illustrating the rescue experiment in *PSTPIP2*-cKI mice. **(B)** Fluorescence-labelled AAV9–*PSTPIP2* was localized in the kidney. **(C)** Real-time PCR analysis of *PSTPIP2* in kidney tissues (n = 6 per group). **(D)** Immunofluorescence (IF) of PSTPIP2 in kidney tissues (n = 6 per group). Scale bar, 50 μm. **(E)** Western blotting analyses of PSTPIP2 (n = 6 per group). PSTPIP2 levels were successfully restored in *PSTPIP2*-cKI mice. **(F and G)** Serum blood urea nitrogen (BUN) and creatinine (Cr) assay. Serum BUN and Cr levels showed that silencing *PSTPIP2* can aggravate kidney damage in *PSTPIP2*-cKI mice treated with AAI (n = 6 per group). **(H)** Representative H&E-stained images of kidney tissues (n = 6 per group). Scale bar, 50 μm. **(I)** Western blotting analysis of KIM-1. Quantification of the KIM-1/β-actin ratio (n = 6). **(J)** Immunohistochemistry (IHC) and kidney injury molecule-1 (KIM-1) levels in kidney tissues (n = 6 per group). Scale bar, 50 μm. **(K)** TUNEL staining and analysis of kidney sections (n = 6 per group). Scale bar, 50 μm. The results showed that silencing *PSTPIP2* can aggravate renal apoptosis in *PSTPIP2*-cKI mice treated with AAI. **(L)** Detection of caspase-3 activity in kidney tissues (n = 6 per group). **(M)** Western blot analysis of cleaved caspase-3. Quantification of the cleaved caspase-3/β-actin ratio (n = 6). **(N)** IHC and cleaved caspase-3 levels in kidney tissues. Scale bar, 50 μm (n = 6 per group). Data are presented as the mean ± SEM of six biological replicates per condition. Each dot represents a sample. Statistically significant differences were determined by an independent sample *t*-test. *P < 0.05, **P < 0.01, ***P < 0.001, ns: non-significant. **Figure 3—source data 1** **Figure 3—source data 2** **AAV9-mediated *PSTPIP2* restoration re-induces renal injury and kidney apoptosis in *PSTPIP2*-cKI mice.**

### Changes in renal *PSTPIP2* expression affects neutrophil infiltration in AAI induced acute renal injury

Inflammation, characterized by neutrophil infiltration, is concomitant with tubular injury and universally presumed to be implicated in the development of AKI. We detected Ly6G as a neutrophil marker, and observed a prominent accumulation of Ly6G^+^ neutrophils in the kidneys of AAI-treated mice (Figure 4A). The infiltrates were accompanied by upregulation of pro-inflammatory TNF-α and chemotactic MCP-1 at the mRNA and protein levels (Figure 4B–E). We found that the infiltration of Ly6G^+^ neutrophils (Figure 4F), as well as the mRNA and protein levels of TNF-α and MCP-1 (Figure 4G–J), were significantly reduced in AAI-treated *PSTPIP2*-cKI mice compared to those in FF mice. IHC staining, ELISA, and real-time PCR analyses showed that restoration of PSTPIP2 expression induced neutrophil infiltration and inflammatory cytokine production in AAI-induced *PSTPIP2*-cKI mice (Figure 4K–O). Hence, we concluded that PSTPIP2 may be involved in neutrophilinfiltration and cytokine production, which was reduced by renal PSTPIP2 overexpression.

**Figure 4.**
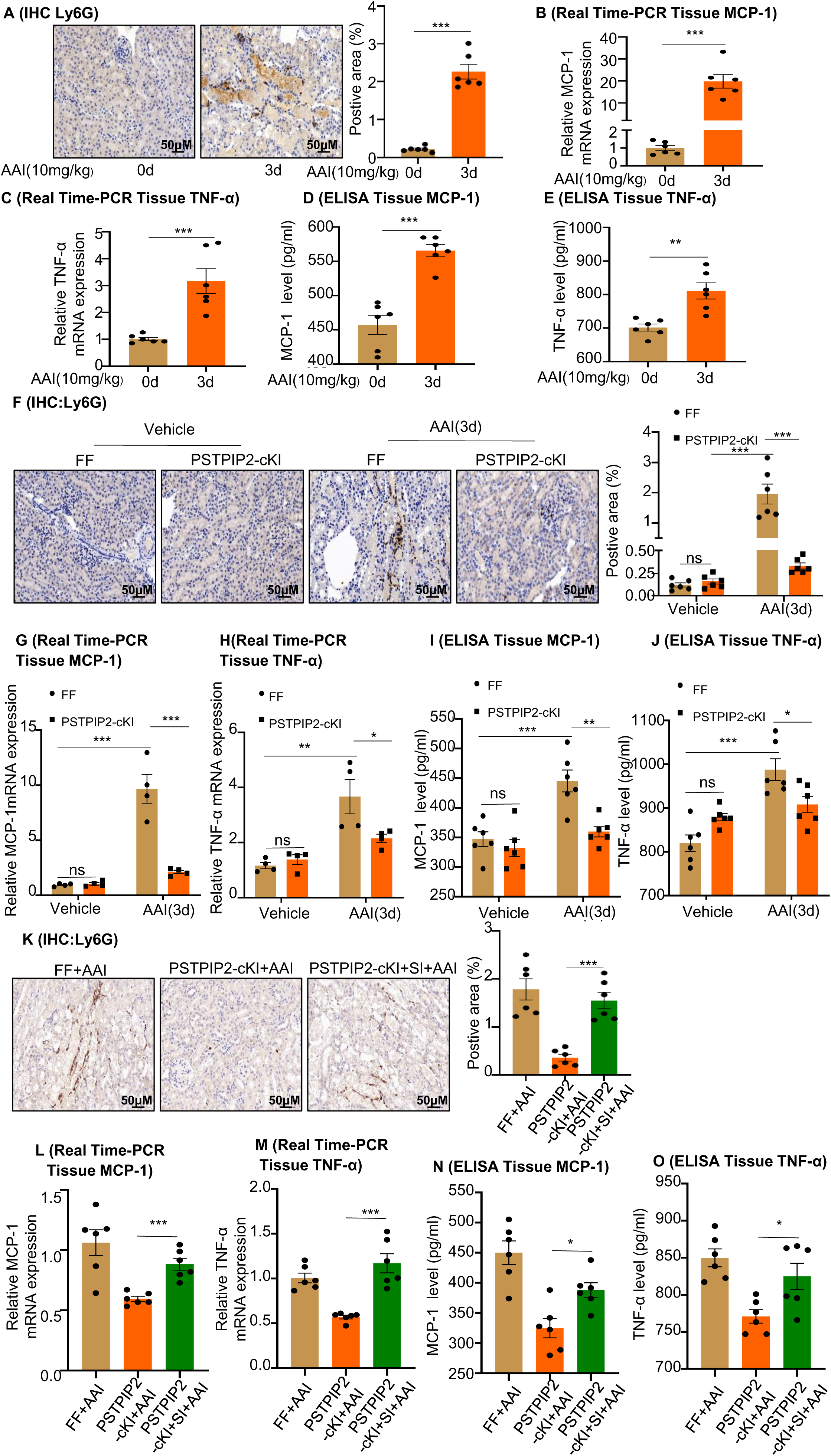
Changes in renal *PSTPIP2* expression affect neutrophil infiltration and inflammatory factor production in mice with aristolochic acid I (AAI)-induced acute renal injury. **(A)** Representative images of immunohistochemistry (IHC) staining with Ly6G^+^. Scale bar, 50 μm. Quantification of the percentage of Ly6G^+^ neutrophil infiltration. n = 6. **(B and C)** mRNA level of renal tissue monocyte chemoattractant protein-1 (*MCP-1*) and tumor necrosis factor-α (*TNF-α*) assessed by real-time PCR (n = 6 per group). **(D and E)** Protein level of renal tissue MCP-1 and TNF-α assessed by ELISA (n = 6 per group). **(F)** Representative IHC staining images of kidney sections, showing the expression of Ly6G. Kidneys were isolated from *PSTPIP2*^Flox/Flox^ and *PSTPIP2*-cKI mice on day 3 after AAI treatment. Scale bar, 50 μm. Quantification of the percentage of Ly6G^+^ neutrophil infiltration (n = 6 per group). **(G and H)** Relative mRNA levels of *MCP-1* and *TNF-α* in renal tissue determined by real-time PCR (n = 4 per group). **(I and J)** Protein level of renal tissue MCP-1 and TNF-α assessed by ELISA (n = 6 per group). The results showed that overexpressing *PSTPIP2* in the kidneys can reduce renal inflammation in mice treated with AAI. **(K)** Representative IHC staining images of kidney sections, showing the expression of Ly6G. Kidneys were isolated from *PSTPIP2*^Flox/Flox^, *PSTPIP2*-cKI, and *PSTPIP2*-cKI-silenced mice on day 3 after AAI treatment. Scale bar, 50 μm. Quantification of the percentage of Ly6G^+^ neutrophil infiltration (n = 6 per group). (**L and M**) Relative mRNA levels of *MCP-1* and *TNF-α* in renal tissue determined by real-time PCR (n = 6 per group). **(N and O)** Protein level of renal tissue MCP-1 and TNF-α assessed by ELISA (n = 6 per group). The results showed that silencing *PSTPIP2* can aggravate renal inflammation in *PSTPIP2*-cKI mice treated with AAI (n = 6 per group). Data are presented as the mean ± SEM of six biological replicates per condition. Each dot represents a sample. Statistically significant differences were determined by an independent sample *t*-test (B–E and K–O) and one-way ANOVA followed by Tukey’s post hoc test (F–J). *P < 0.05, **P < 0.01, ***P < 0.001, ns: non significant. **Figure 4—source data 1** **Changes in renal *PSTPIP2* expression affect neutrophil infiltration and inflammatory factor production in mice with aristolochic acid I (AAI)-induced acute renal injury.**

We hypothesized that the transient depletion of neutrophils may provide therapeutic benefits in AAI-induced kidney injury. We first examined whether treatment with a specific Ly6G neutralizing antibody 1 d before and after AAI injection could deplete their circulating and renal neutrophil levels. Control mice were treated with the same dosage of rat IgG isotype control antibody (Figure S3A). Compared to the control, we found that the Ly6G antibody could specifically prevent the AAI-induced increase in circulating and renal neutrophil numbers (Figure S3B–D). Next, we assessed the *in vivo* therapeutic benefits of neutrophil depletion in AAN. Neutrophil depletion significantly reduced the levels of serum Cr and BUN in AAI-treated mice compared to those in control antibody-treated mice (Figure S4A and B). In addition, neutrophil depletion in AAI-treated mice significantly attenuated tissue damage and downregulated KIM-1 expression, based on H&E staining and IHC analysis, respectively (Figure S4C, D), in Ly6G antibody-treated mice. Neutrophil depletion prevented renal dysfunction, highlighting the integral contribution of neutrophils to AAN. Furthermore, we found that neutrophil depletion significantly decreased TUNEL^+^ cell numbers (Figure S4E) and significantly downregulated the activity of caspase-3 and level of cleaved caspase-3 following neutrophil depletion in Ly6G antibody-treated mice (Figure S4F and G). These results suggest that neutrophil depletion ameliorates AAI-induced kidney damage and apoptosis in RTECs.

### Changes in *PSTPIP2* expression affects NET formation in AAI-induced acute renal injury

Although increased neutrophil infiltration in the kidney is the major cause of AAN, the mechanism of NET formation following AAI exposure remains unknown. To evaluate NET formation, mice received a single IP injection of AAI and were sacrificed after 3 d. Serum nucleosome, MPO, and dsDNA levels were significantly increased after AAI treatment (Figure 5A–C). Cit-H3 protein levels, another marker of NETs, were also significantly increased after AAI treatment, as observed by fluorescence microscopy and western blotting (Figure 5D and E). We found that serum nucleosome, MPO, and dsDNA levels (Figure 5F–H), as well as Cit-H3 levels (Figure 5I and J), were significantly lower in AAI-treated *PSTPIP2*-cKI mice than those in AAI-treated FF mice. Finally, we found that serum levels of nucleosomes, dsDNA, and MPO, as well as the expression of Cit-H3, were significantly increased in AAI-treated mice after PSTPIP2 expression was restored compared with those in AAI-treated *PSTPIP2*-cKI mice (Figure S5A–E). Our results demonstrate that PSTPIP2 may play a vital role in the regulation of NET formation.

**Figure 5.**
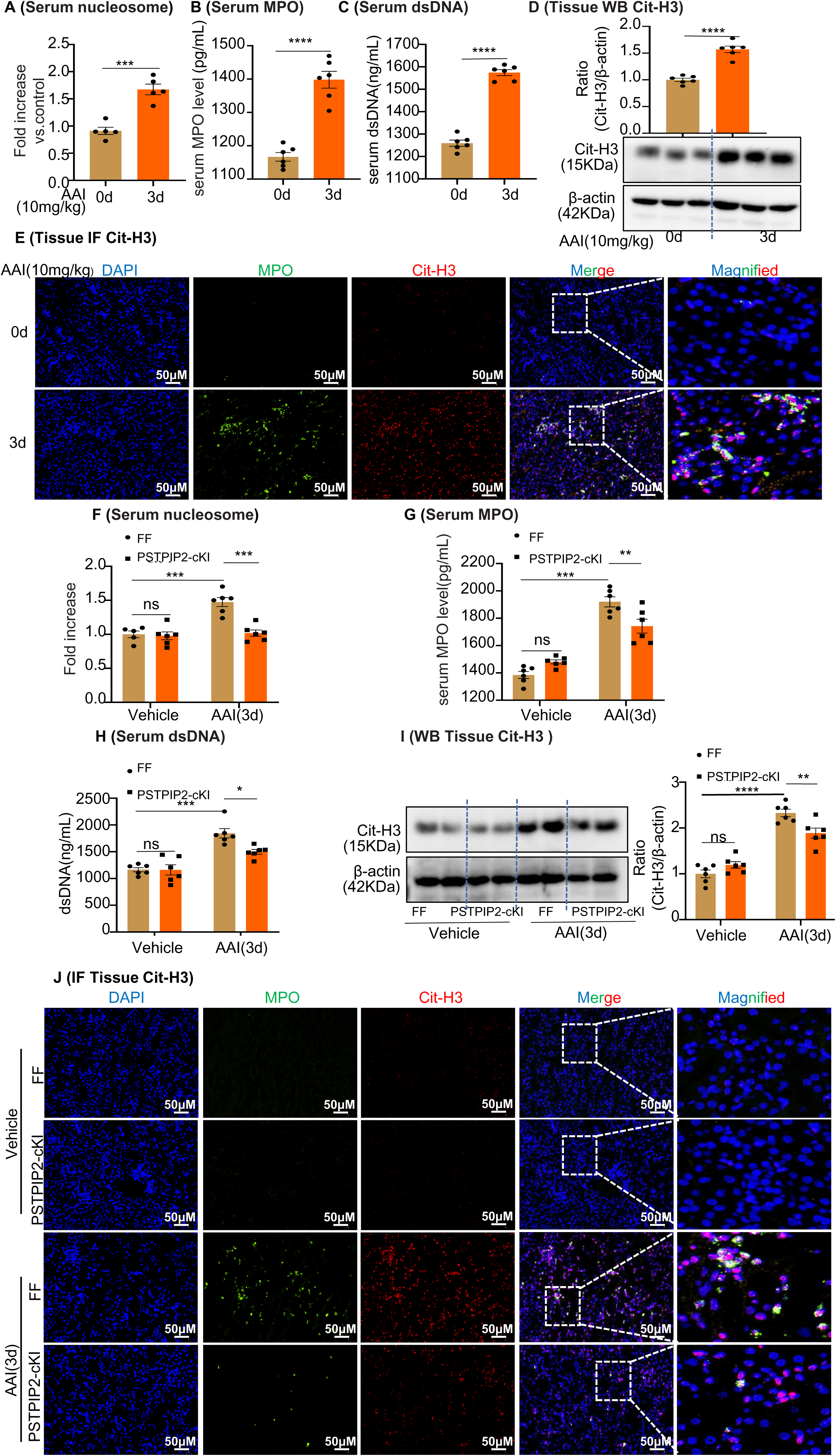
*PSTPIP2* conditional knock-in in the kidney significantly decreased neutrophil extracellular trap (NET) formation. **(A)** Serum nucleosome level after aristolochic acid I (AAI) treatment for 3 d (n = 5). **(B and C)** Serum myeloperoxidase (MPO) and dsDNA level after AAI treatment for 3 d (n = 6). **(D)** Protein level of citrullinated histone 3 (Cit-H3) assessed by western blotting after AAI treatment for 3 d (n = 6). **(E)** Immunofluorescent (IF) staining revealed increased co-localization of MPO (green) and Cit-H3 (red) after AAI treatment for 3 d (n = 6). Scale bar, 50 μm. **(F)** Serum nucleosome level of *PSTPIP2*^Flox/Flox^ and *PSTPIP2*-cKI mice treated with AAI (n = 6 per group). **(G and H)** Serum MPO and dsDNA level of *PSTPIP2*^Flox/Flox^ and *PSTPIP2*-cKI mice treated with AAI (n = 6 per group). **(I)** Western blot analysis of Cit-H3. Quantification of the Cit-H3/β-actin ratio (n = 6 per group). **(J)** Representative IF staining images of kidney sections, showing the expression of Cit-H3 (red) and MPO (green). Kidneys were isolated from *PSTPIP2*^Flox/Flox^ and PSTPIP2-cKI mice on day 3 after AAI treatment (n = 6 per group). Scale bar, 50 μm. Data are presented as the mean ± SEM of six biological replicates per condition. Each dot represents a sample. Statistically significant differences were determined by an independent sample *t*-test (A–D) and one-way ANOVA followed by Tukey’s post hoc test (F–I). *P < 0.05, **P < 0.01, ***P < 0.001, ****P < 0.0001, ns: non-significant. **Figure 5—source data 1** **Figure 5—source data 2** ***PSTPIP2* conditional knock-in in the kidney significantly decreased neutrophil extracellular trap (NET) formation.**

NET formation during programmed neutrophil cell death (NETosis) has recently been shown to play a pro-injury role in renal inflammatory diseases (18). Accordingly, we explored whether NETosis increases kidney injury induced by AAI. For this purpose, we intravenously injected DNase I or GSK484, a potent inhibitor of NET formation, into the AAI-treated mice (Figure S6A). Serum nucleosome, MPO, and dsDNA levels (Figure S6B–D), as well as Cit-H3 levels (Figure S6E and F), were significantly decreased in mice treated with DNase I or GSK484. We tested the effects of DNase I and GSK484 on the progression of AAN. Treatment with DNase I or GSK484 significantly reduced serum Cr and BUN levels (Figure S7A and B) and attenuated tissue damage (Figure S7C) in AAI-treated mice. Moreover, IHC analysis showed that the KIM-1 and TUNEL^+^ cell levels were significantly decreased in DNase I- or GSK484-treated mice (Figure S7D and E). The activity of caspase-3 and level of cleaved caspase-3 protein were also significantly downregulated in AAI-treated mice treated with GSK484 or DNase I (Figure S7F and G). Based on these observations, we surmised that the capacity of neutrophils to cause or exacerbate apoptosis is linked to their NET formation properties. Primary renal neutrophils were isolated from AAI induced DNase I-treated mice (Figure S8A) and the levels of dsDNA and neutrophil elastase in their supernatants were examined. We found that dsDNA and neutrophil elastase production could be inhibited by DNase I (Figure S8B–C). Similarly, Cit-H3 protein levels were significantly decreased in primary renal neutrophils of AAI induced DNase I-treated mice (Figure S8D and E). To examine the toxicity of AAI induced NETs on tubular cells, RTECs were exposed to the supernatant of NETs (Figure S8F). After 20 h of incubation, we found decreased numbers of apoptotic cells, as well as downregulated caspase-3 activity and cleaved caspase-3 levels, after DNase I and AAI treatments (Figure S8G–J). These results suggest that inhibition of NET formation attenuates AAI-induced kidney damage.

### *PSTPIP2* inhibits NF-κB signaling to reduce the release of IL-19 from damaged RTECs, thereby reducing neutrophil infiltration and NET formation

To better characterize the specific role of PSTPIP2 in RTECs, overexpression of PSTPIP2 was induced by transfection with the PSTPIP2 plasmid (pEX-2-PSTPIP2) in mRTECs. We confirmed that the pEX-2-PSTPIP2 plasmid elevated PSTPIP2 expression (Figure S9D–E). We stimulated RTECs with AAI for 20 h to explore whether they secrete important cytokines (Figure 6A). Real-time PCR analysis showed that the expression of IL-19 was most significantly elevated compared with other interleukins in AAI-treated mRTECs, and overexpression of PSTPIP2 significantly decreased IL-19 expression compared with that in control plasmid transfected cells (Figure 6B). We found that the level of IL-19 was significantly increased in the supernatant of AAI-treated mRTECs (Figure 6C), consistent with a drastic increase in its mRNA and protein levels (Figure 6D and E). Conversely, we found that the expression of IL-19 was significantly reduced after overexpression of PSTPIP2 in mRTECs (Figures 6K-L and S9F). Using immunofluorescence staining, we observed that E-cadherin was colocalized with IL-19, suggesting that RTECs are the major source of IL-19 after AAI-induced renal injury (Figure S9A). Serum IL-19 levels have been shown to be increased with various kidney diseases, including kidney transplants, hydronephrosis accompanied by ureteral stones, diabetes, and polycystic kidney disease (Table S1). Notably, serum IL-19 and Cr levels were positively correlated (*R* = 0.5472, P < 0.05; Figure 6F). The expression of IL-19 was significantly lower in AAI-treated *PSTPIP2*-cKI mice than that in AAI-treated FF mice (Figures 6G,I and S9B). However, the restoration of PSTPIP2 expression significantly increased IL-19 expression after stimulation with AAI (Figures 6H,J and S9C).

**Figure 6.**
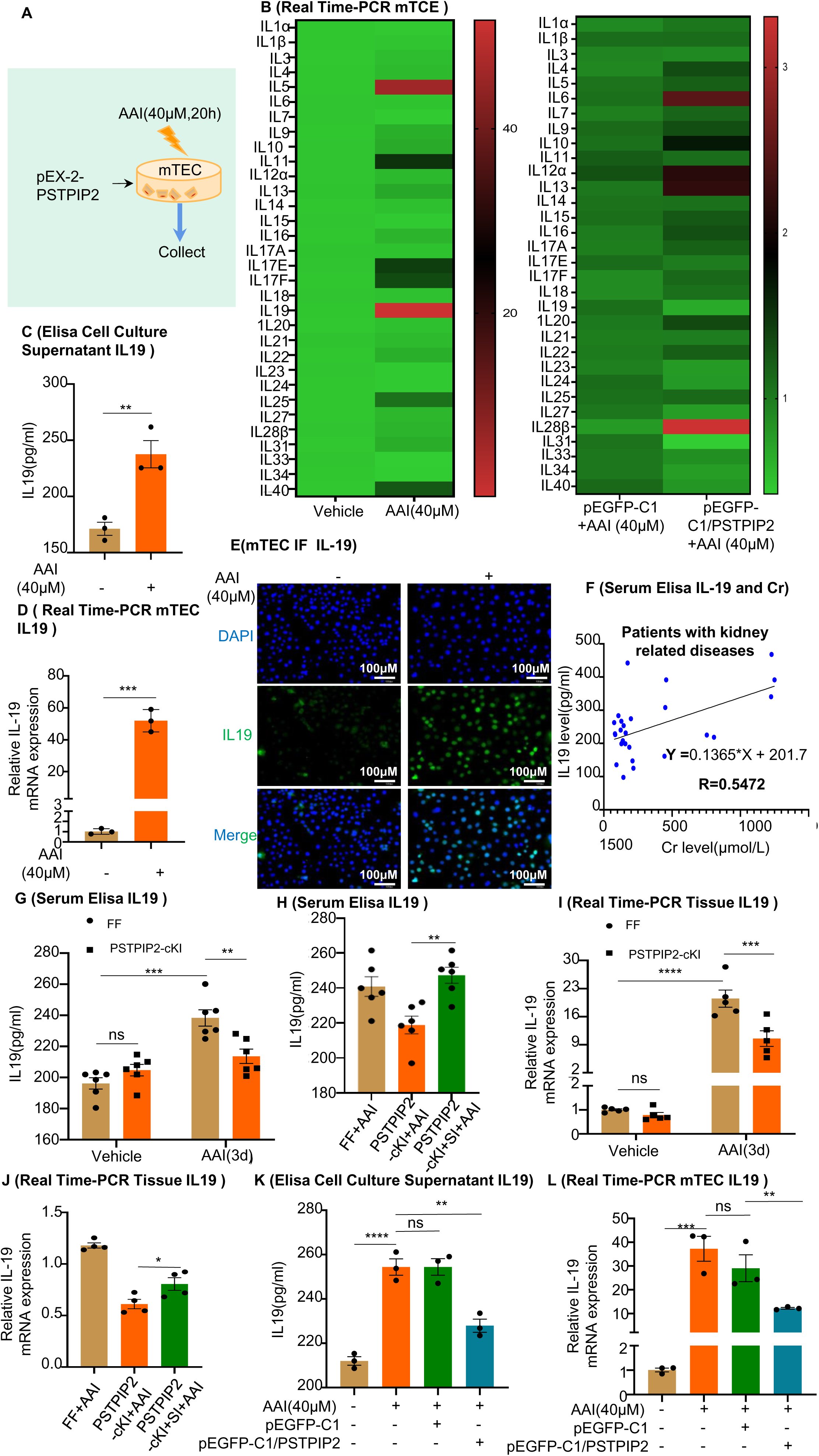
Expression of interleukin-19 (IL-19) was associated with kidney injury and regulated by *PSTPIP2*. **(A)***PSTPIP2* was overexpressed in mouse renal tubular epithelial cells (mRTECs) by the plasmid pEGFP-C1/*PSTPIP2*; the cells were subsequently treated with aristolochic acid I (AAI, 40 μM) for 20 h. **(B)** mRNA levels of various interleukins in mRTECs assessed by real-time PCR (n = 3). **(C)** Protein level of IL-19 in the cell culture supernatant assessed by ELISA (n = 3). **(D)** mRNA level of *IL-19* in mRTECs assessed after treatment with AAI by real-time PCR (n = 3). **(E)** IL-19 levels were assessed by immunofluorescence (IF) staining in cells treated with AAI. Scale bar, 100 μm (n = 3). **(F)** Correlation analysis of serum IL-19 and Cr levels in patients with kidney-related diseases (n = 27). **(G)** Serum levels of IL-19 protein in *PSTPIP2*^Flox/Flox^ and *PSTPIP2*-cKI mice treated with AAI (n = 6). **(H)** Serum levels of IL-19 protein in *PSTPIP2*^Flox/Flox^, *PSTPIP2*-cKI, and *PSTPIP2*-cKI silenced mice on day 3 after AAI treatment (n = 6). **(I)** Relative mRNA levels of IL-19 in *PSTPIP2*^Flox/Flox^ and *PSTPIP2*-cKI mice treated with AAI determined by real time PCR (n = 5 per group). **(J)** Real-time PCR analysis of IL-19 (n = 4 per group). **(K)** Protein level of IL-19 in cell culture supernatant assessed by ELISA (n = 3). **(L)** mRNA level of *IL-19* in mRTECs assessed by real-time PCR (n = 3). Data are presented as the mean ± SEM of three to six biological replicates per condition. Each dot represents a sample. Statistically significant differences were determined by an independent sample *t*-test (B-D,H,J) and one-way ANOVA followed by Tukey’s post hoc test (G,I,K,L). Data in F were analyzed by Pearson correlation.*P < 0.05, **P < 0.01, ***P < 0.001, ****P < 0.0001, ns: non-significant. **Figure 6—source data 1** **Expression of interleukin-19 (IL-19) was associated with kidney injury and regulated by *PSTPIP2*.**

IL-19 is a potent cytokine in neutropenia treatment (19). We assessed whether IL-19 is chemotactic to mouse bone marrow-derived neutrophils using a Transwell system (Figure 7A) and found that IL-19 stimulation was able to induce neutrophil migration, especially at a concentration of 80 ng/mL (Figure 7B). Although IL-19 has been shown to enhance neutrophil recruitment, it is unclear whether IL-19 induces NET formation. We used a series of different concentrations of recombinant IL-19 (rIL-19) to stimulate mouse bone marrow-derived neutrophils (Figure S9G) for 4 h (Figure 7C). The protein levels of IL-20Rβ, the IL-19 membrane receptor, increased drastically after stimulation with various rIL-19 concentrations (Figure 7G and 7I), suggesting that neutrophils express IL-20Rβ, which is upregulated in response to rIL-19 exposure. We sought to determine whether IL-19 induces NET formation. Accordingly, we measured nucleosome, MPO, and dsDNA levels in neutrophil media after stimulation with rIL-19 and found a significant increase in all levels at a concentration of 80 ng/mL compared to those in the vehicle group (Figure 7D–F). Cit-H3 protein levels were significantly decreased in IL-19-treated mouse bone marrow-derived neutrophils (Figure 7G and 7H). Under similar conditions, the neutrophils lost their nuclei and released strings and cellular debris after IL-19 treatment, as observed with SEM (Figure 7J). To examine the toxicity of IL-19-induced NETs on damaged tubular cells, RTECs were exposed to the supernatant of NETs (Figure S10A); at 20-h of incubation, we found an increased number of apoptotic RTECs (Figure S10D and E). Consistently, the activity of caspase-3 and level of cleaved caspase-3 protein were significantly upregulated after stimulation with IL-19-derived NETs (Figure S10B and C).

**Figure 7.**
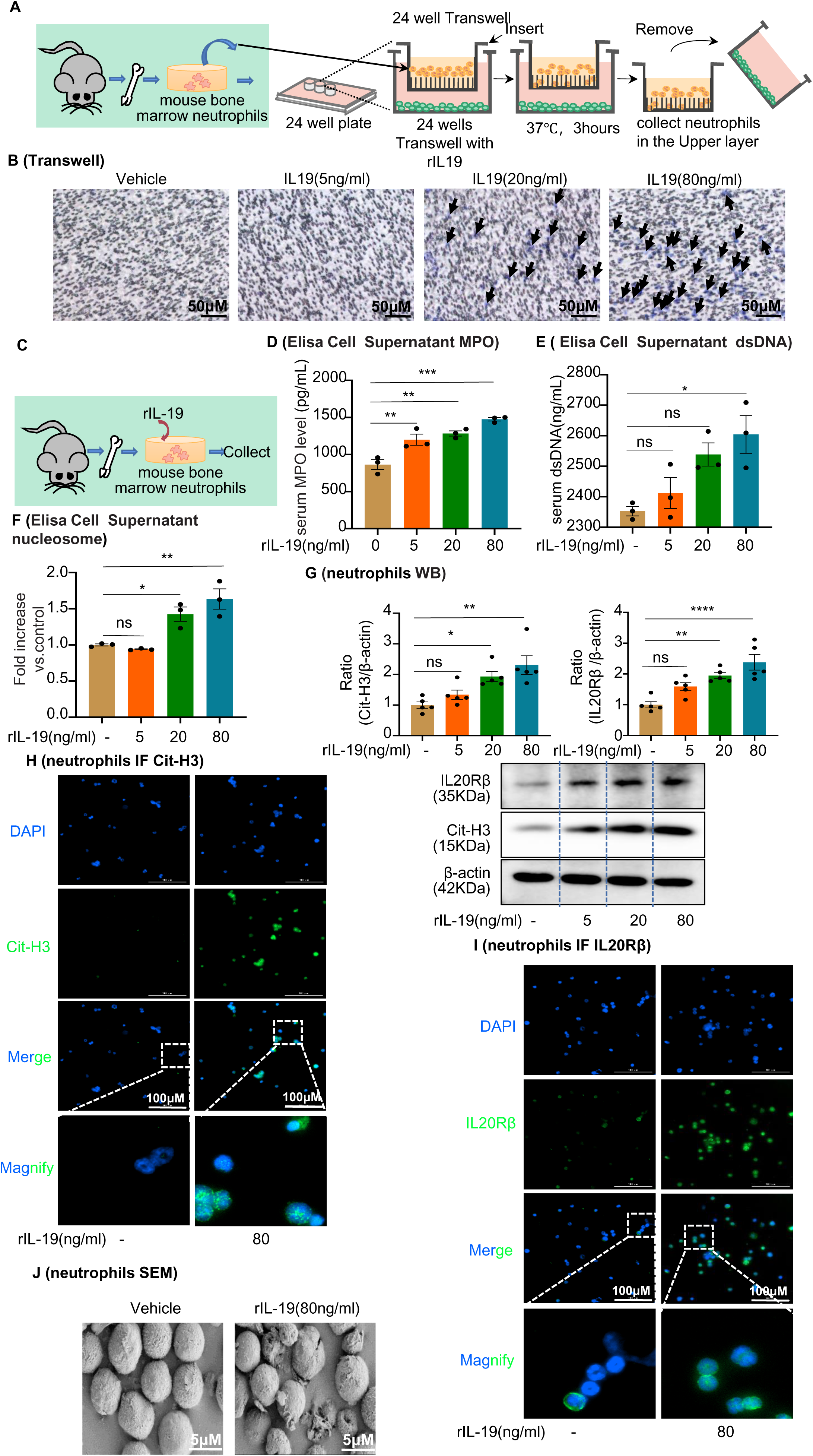
Interleukin-19 (IL-19) induces neutrophil extracellular trap (NET) formation *in vitro* through IL-20Rβ receptor signaling. **(A)** Schematic of Transwell migration assay for neutrophils. **(B)** Neutrophils were isolated from mouse bone marrow and loaded into the upper Transwell chamber. Serum-free RPMI-1640 medium (500 μL) containing 1% penicillin/streptomycin (P/S), with or without different concentrations of IL-19, was added to the lower chamber. Neutrophil migration was detected by crystal violet staining. Neutrophils are marked by black arrows (n = 3). Scale bar, 100 μm. **(C)** Mouse bone marrow-derived neutrophils were treated with different concentrations of IL-19 (5 ng/mL, 20 ng/mL, 80 ng/mL) for 4 h and then collected. **(D and E)** Level of myeloperoxidase (MPO) and dsDNA in the cell culture supernatant assessed after treatment with different concentrations of IL-19 (n = 3). **(F)** Level of nucleosomes in the cell culture supernatant assessed after treatment with different concentrations of IL-19 (n = 3). **(G)** Protein levels of IL-20Rβ and citrullinated histone 3 (Cit-H3) assessed by western blotting after treatment with different concentrations of IL-19 (n = 5). **(H and I)** Immunofluorescence staining of IL-20Rβ and Cit-H3 of neutrophils after treatment with different concentrations of IL-19 (n = 3). Scale bar, 100 μm. **(J)** Representative scanning electron microscopy images of vehicle- and IL-19-stimulated neutrophils (NETs, n = 3). Scale bar, 5 μm. Data are presented as the mean ± SEM of three biological replicates per condition. Each dot represents a sample. Statistically significant differences were determined by an independent sample *t*-test and one-way ANOVA followed by Tukey’s post hoc test (D–G). *P < 0.05, **P < 0.01, ***P < 0.001, ****P < 0.0001, ns: non-significant. **Figure 7—source data 1** **Figure 7—source data 2** **Interleukin-19 (IL-19) induces neutrophil extracellular trap (NET) formation *in vitro* through IL-20Rβ receptor signaling.**

The transcription factor NF-κB has been reported to regulate IL-19 gene transcription (19), but it is unclear whether PSTPIP2 regulates IL-19 expression by activating or inhibiting the NF-κB pathway. In addition, to verify the correlation between PSTPIP2 and NF-κB p65, we used mRTEC extract for co-immunoprecipitation (Co-IP) experiments using anti-NF-κB p65 and anti-PSTPIP2 antibodies. The Co-IP results confirmed an interaction between PSTPIP2 and NF-κB p65 in mRTECs (Figure 8A). As expected, NF-κB was strongly activated in AAI-treated mRTECs, as evidenced by the markedly increased phosphorylation (S536) and nuclear localization of NF-κB p65, accompanied by enhanced phosphorylation of IκBα (S32/36), and overexpression of PSTPIP2 inhibited AAI-induced activation of NF-κB signaling (Figure 8B and C). In addition, single IF staining showed that the overexpression of PSTPIP2 significantly inhibited the nuclear transfer of p65 (Figure 8D). The NF-κB pathway was inhibited after a 2-h pretreatment with the NF-κB inhibitor PDTC (25 μM; Figure 8F), immediately after which, we removed the cell supernatant and retreated the cells with AAI for 20 h (Figure 8E). Real-time PCR analysis showed that IL-19 was visibly upregulated in the PDTC+AAI-treated group compared to the AAI treated group (Figure 8G). Collectively, these results indicate that PSTPIP2 inhibits NF-κB signaling to induce IL-19 release from the renal tubular epithelial cells.

**Figure 8.**
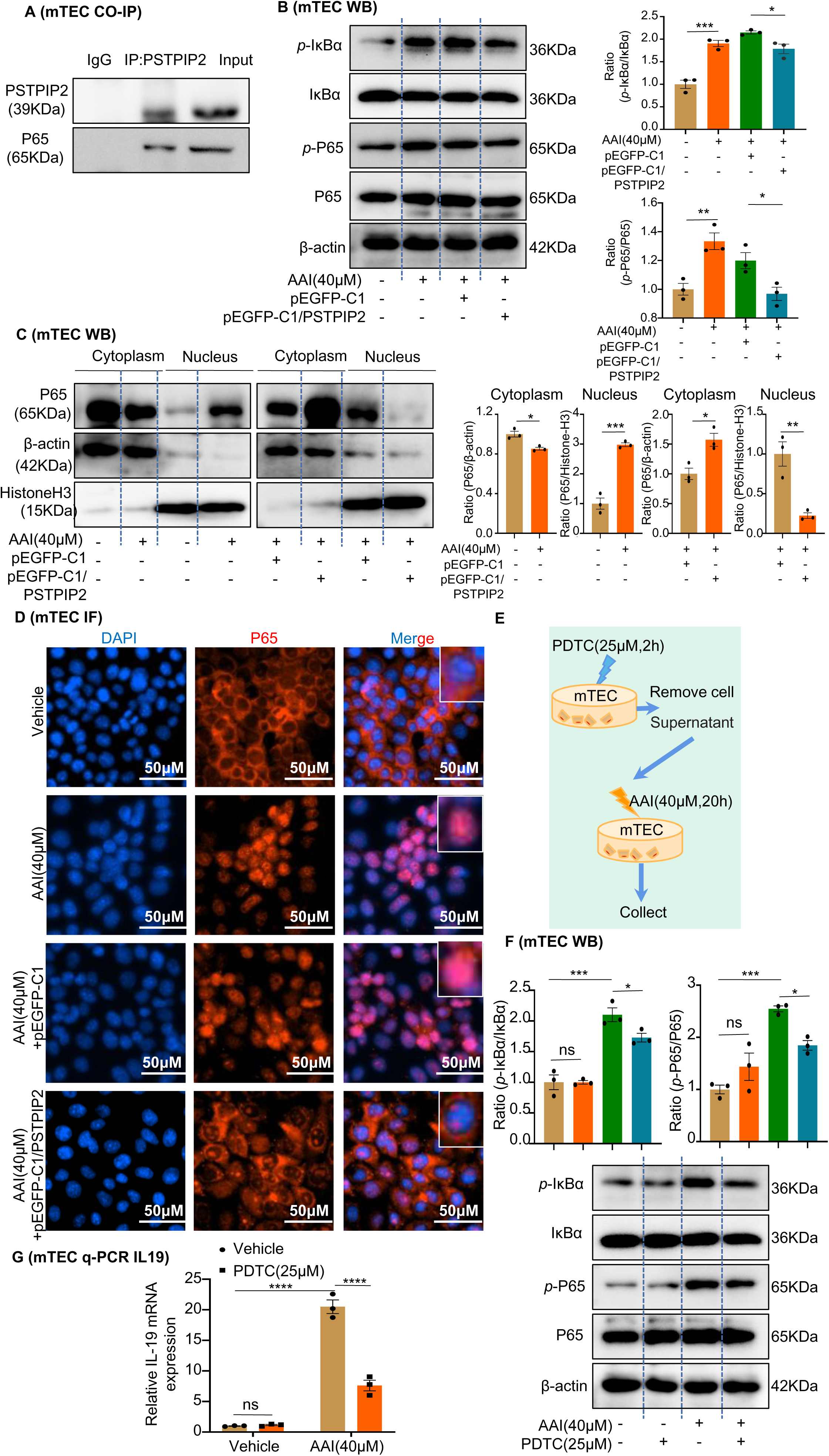
*PSTPIP2* reduces interleukin-19 (IL-19) transcription by promotion of NF-κB signaling. **(A)** Co-immunoprecipitation (Co-IP) of *PSTPIP2* and NF-κB p65 in mRTEC using anti-NF-κB p65 and anti-*PSTPIP2* antibodies. **(B)** Western blotting results showed that the protein expression of phosphorylated IκBα and p65 was significantly increased after AAI stimulation. Overexpression of *PSTPIP2* significantly reduced AAI-induced upregulation of phosphorylated IκBα and p65 proteins (n = 3). (**C**) Expression of NF-κB p65 in the nucleus and cytoplasm of various groups of mRTECs; β-actin and histone H3 were used as cytoplasmic and nuclear protein markers, respectively (n = 3). **(D)** Immunofluorescence was used to detect the effect of *PSTPIP2* overexpression on the nuclear transfer of p65 (n = 3). **(E)** NF-κB signaling was inhibited after a 2-h pretreatment with the NF-κB inhibitor PDTC (25 μM, n = 3). **(F)** Western blot analysis of phosphorylated IκBα and p65 (n = 3). **(G)** mRNA level of *IL-19* in AAI-treated mRTECs exposed to the NF-κB inhibitor PDTC (25 µM, 2 h) measured by qPCR (n = 3). Data are presented as the mean ± SEM of three biological replicates per condition. Each dot represents a sample. Statistically significant differences were determined by an independent sample *t*-test and one-way ANOVA followed by Tukey’s post hoc test (B,C,F,G). *P < 0.05, **P < 0.01, ***P < 0.001, ****P < 0.0001, ns: non-significant. **Figure 8—source data 1** **Figure 8—source data 2** ***PSTPIP2* reduces interleukin-19 (IL-19) transcription by promotion of NF-κB signaling.**

## Discussion

This study provides a new perspective on the role of PSTPIP2 in acute AAN and identifies a potential mechanism of PSTPIP2 in inhibiting epithelial apoptosis to alleviate AAI-induced nephrotoxicity. We confirmed that PSTPIP2 was significantly downregulated in both AAI-induced acute AAN mouse models and AAI-exposed mRTECs. We also found that the conditional knock-in of PSTPIP2 in mouse kidneys was sufficient to ameliorate renal dysfunction, histological injury, cell apoptosis, and inflammatory responses induced by AAI in acute AAN mouse models. Accordingly, the restoration of normal PSTPIP2 expression in *PSTPIP2*-cKI mice promoted renal injury and cell apoptosis, confirming the critical role of PSTPIP2 in acute AAN. Mechanistically, we found that PSTPIP2 served as a negative regulator of acute AAN, at least in part, via reducing apoptosis of RTECs by regulating the IL-19-mediated formation of NETs. Our results therefore support the potential of PSTPIP2-targeted therapy in patients with acute AAN.

First, we examined the role of PSTPIP2 in renal injury and inflammation. PSTPIP2 plays a crucial role in autoimmune diseases and is involved in macrophage activation, neutrophil migration, and cytokine production (20). Studies have shown that the protein tyrosine phosphatase PTP-PEST, Src homology domain-containing inositol 5′- phosphatase 1 (SHIP1), and C-terminal Src kinase (CSK) can bind to PSTPIP2 and inhibit the development of autoinflammatory diseases (21). Yang *et al*. (22) demonstrated that PSTPIP2 could ameliorate the degree of liver fibrosis and hepatic inflammation in CCl_4_-induced hepatic fibrosis. Neutrophils represent the first-line of innate immune defense to clear or contain invading pathogens. In AAN, which is characterized by increased inflammation and cell death (23), an increase in renal neutrophil levels was associated with poor clinical outcomes (24). Recent studies have also shown that increased neutrophil levels can exacerbate AKI (25, 26). We observed a large infiltration of neutrophils in the kidneys and a significant increase in the expression of pro-inflammatory TNF-α and chemotactic MCP-1 molecules 3 d after establishing acute AAN. However, conditional knock-in of PSTPIP2 in the kidney markedly reduced neutrophil infiltration and the expression of inflammatory factors, both of which were significantly increased after restoring PSTPIP2 to normal levels in the kidney. Given the important role of neutrophils in the pathomechanism of AKI, we assessed whether transient neutrophil depletion would induce similar therapeutic benefits in AAN. We found that a Ly6G-specific monoclonal antibody (27) could deplete both circulating and renal neutrophil levels in mice, reducing systemic inflammation and renal injury. Given that neutrophils are an important immune cell type, transient neutrophil depletion may be feasible in the treatment of AAN.

Over the past decade, many new theories regarding the pathogenesis of AAN have emerged (28–30). However, the pathophysiological mechanisms by which AA induces renal injury remain largely unknown. Neutrophils are rapidly recruited to tissues during infections to eliminate pathogens, both by phagocytosis and the formation of NETs in the extracellular space (31). NETs are one of the most widely studied pathways of neutrophil-induced injury and have been observed in kidney injuries induced by various stimuli (32). Thus, there is a growing interest in the synthesis and regulation of NETs. NET formation is usually associated with neutrophil death, or NETosis, which is morphologically distinct from apoptosis and necrosis (33). NET formation depends on the activation of peptidyl arginine deiminase (PAD) enzymes that convert arginine residues of histones to citrulline (34). Histone citrullination neutralizes DNA–histone interactions, resulting in chromatin decondensation and NET release (35). NETs are composed of neutrophil elastase, citrullinated histones, and extracellular DNA (36). Because PSTPIP2 is closely involved in neutrophil migration, we hypothesized that it participates in the formation and regulation of NETs. In our current study, the levels of NET markers, such as nucleosome, MPO, dsDNA, and Cit-H3, were significantly increased in AAI-exposed mice, and conditional knock-in of PSTPIP2 expression in the kidney reduced NET formation. However, reducing PSTPIP2 expression in mouse kidneys significantly increased NET formation. Indeed, a previous study showed that AAN kidneys were protected by treatment with the inhibitor of NET formation GSK484 or DNase I (37), which is consistent with the findings in glomerular disease (38).

PSTPIP2 plays an important role in cytokine production. Notably, the expression of IL-19 was the most significantly elevated among the interleukins in AAI-treated mRTECs. Overexpression of PSTPIP2 significantly decreased *IL-19* mRNA expression. In the current study, we performed a complementary retrospective association study using serum samples from patients with various kidney diseases. The results indicated that IL-19 was positively correlated with serum Cr level (Figure 6F), a classic indicator of kidney injury. IL-19 is a member of the IL-10 family of cytokines, which has indispensable functions in many inflammatory processes. It is also a member of the IL-20 subfamily and binds to the IL-20 receptor to regulate various processes, such as antimicrobial activity, wound healing, and tissue remodeling (39). Cellular sources have been characterized as having myeloid and epithelial origins, which correspond with the significant levels found in the RPTEC/TERT1 renal epithelial cell system. IL-19 participates in several animal and human diseases, including psoriasis, inflammatory bowel disease, atherogenesis, endotoxic shock, rheumatoid arthritis, and cancer (39). As interleukins play a pivotal role in the formation of NETs, Hamza *et al.* (40) showed that IL-33 is released from liver sinusoidal endothelial cells to promote NET formation during liver ischemia– reperfusion, exacerbating inflammatory cascades and sterile inflammation. Nie *et al.* (41) demonstrated that IL-8-induced NET formation promoted diffuse large B-cell lymphoma progression via TLR9 signaling. IL-19, produced predominantly by osteocytes, stimulates granulopoiesis and neutrophil formation, which in turn stimulates IL-20Rβ/Stat3 signaling in neutrophil progenitors to promote their expansion and neutrophil formation (19). To date, the relationship between IL-19 and NET formation has not been reported. In the present study, we found that IL-19 can chemotax neutrophils and bind to the surface receptor IL-20Rβ to form NETs. Further studies found that PSTPIP2 could affect IL-19 expression by regulating the NF-κB pathway in RTECs. After PSTPIP2 conditional knock-in in mouse kidneys, the expression of IL-19 was inhibited, whereas it was noticeably increased after reducing the expression of PSTPIP2. Therefore, PSTPIP2 affects NET formation precisely because of its regulation of IL-19 release in RTECs.

Finally, it is well accepted that RTEC injury and apoptosis are common histopathological features of AKI (42). Although programmed cell death is necessary to maintain health, its dysregulation by excessive or defective apoptosis has been implicated in a variety of diseases (43). Experimental evidence supports a pathogenic role of apoptosis in AKI. Wei *et al.* (44) reported that inhibition of RTEC apoptosis protected mice from ischemic AKI. We previously found that apoptosis of RTECs contributed to cisplatin nephrotoxicity (45). We also showed for the first time that histone acetylation-mediated silencing of PSTPIP2 may contribute to cisplatin nephrotoxicity. PSTPIP2 may serve as a potential therapeutic target in the prevention of cisplatin nephrotoxicity. Nevertheless, further studies are needed to determine the mechanism of PSTPIP2-mediated suppression of epithelial cell apoptosis (46). NETs are important for immune defense; however, their role in mediating apoptosis remains controversial. In the current study, TUNEL^+^ cell numbers and caspase-3 activity were significantly increased in the kidneys of mice with AAN, whereas they were significantly decreased in the kidneys of PSTPIP2 conditional knock-in mice. Conversely, the level of apoptosis was markedly increased with a reduction in renal PSTPIP2 expression. We investigated whether NETs also developed during AAI induced apoptosis and found that the kidneys of neutrophil-depleted mice treated with the Ly6G antibody and NET-depleted mice treated with GSK484 (a PAD4 inhibitor) or DNase I (a nonspecific inhibitor of NETs) showed a significant decrease in the number of TUNEL^+^ cells. *In vitro*, we showed that conditioned NET media pre-treated with DNase I reduced tubular cell apoptosis. We treated injured tubular epithelial cells by stimulating neutrophil-derived NETs with exogenous IL-19 and found significantly increased levels of apoptosis.

This study investigated the renoprotective effects of PSTPIP2 in AAI-induced AAN and revealed that NETs participated in both AAI-induced renal injury and apoptosis in RTECs. Moreover, PSTPIP2 reduced IL-19 secretion by inhibiting activation of the NF-κB pathway in RTECs, which in turn reduced the generation of NETs (Figure 9).

**Figure 9.**
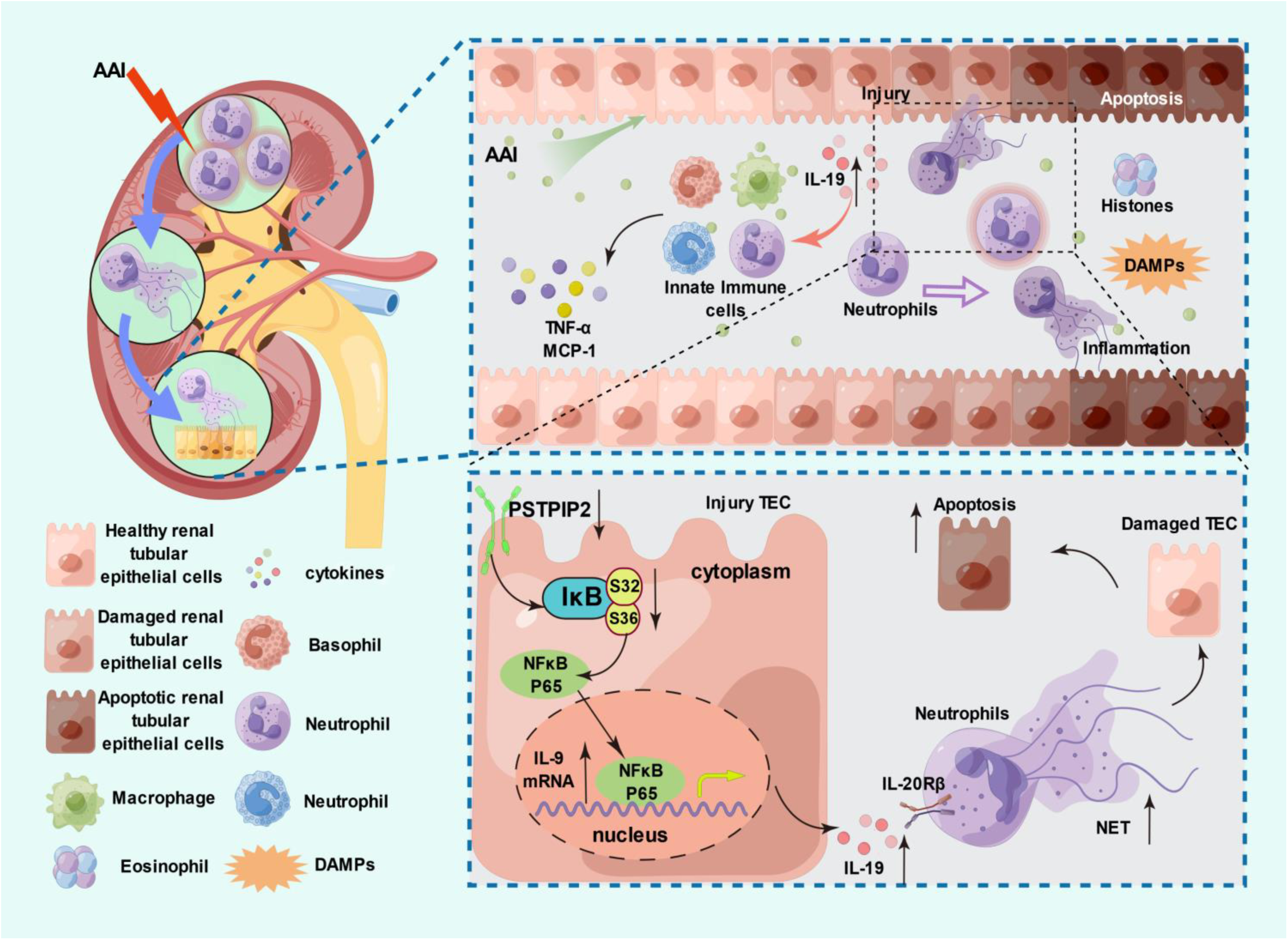
*PSTPIP2* alleviates aristolochic acid-induced acute kidney injury and renal tubular epithelial cell apoptosis by suppressing interleukin-19-mediated neutrophil extracellular trap formation.

## Materials and Methods

### Experimental materials

A complete list of the antibodies, reagents, kits, and other necessary materials used in the experiments can be found in the Supplementary Methods.

### Mouse model of AAN

Six- to eight-week-old male C57BL/6J mice(20-22g) were obtained from the Experimental Animal Center of the Anhui Medical University. All animal experiments were performed in accordance with the Regulations of the Experimental Animal Administration issued by the State Committee of Science and Technology of China. Efforts were made to minimize the number of animals used and their suffering. Animals were maintained in accordance with the guidance of the Center for Developmental Biology, Anhui Medical University, for the care and use of laboratory animals, and all experiments used protocols approved by the institutions’ subcommittees on animal care. (approval No. LLSC20200730). All mice were randomly assigned to a treatment group and housed in a comfortable environment with a 12 h/12 h light/dark cycle for 1 week before experimentation. Mice were administered AAI (10 mg/kg) in a single intraperitoneal (IP) injection, and six mice per group were separately sacrificed at 1-, 3-, and 5-d post injection. Control mice were IP injected with the same dosage of vehicle.

### Neutrophil depletion

Systemic neutrophil depletion was achieved using an IP injection of purified anti mouse Ly6G (10 µg/g) before and 1 d after the AAI injection. Control mice were treated with the same dosage of rat IgG isotype control, and six mice per group were separately sacrificed at 3-d post injection.

### DNase I and GSK484 treatment

DNase I (10 U/100 μL, 0.9% NaCl) or GSK484 (4 mg/kg) was IP injected daily for 3 d prior to AAI treatment, and six mice per group were separately sacrificed at 3-d post-injection.

### Neutrophil isolation

Mouse bone marrow-derived neutrophils were isolated and purified by density gradient centrifugation as previously reported (47, 48). Further information on the extraction of mouse bone marrow- and renal tissue-derived primary neutrophils can be found in the Supplementary Methods.

### NET formation assay

Mouse bone marrow-derived neutrophils were treated with 40 μM AAI or DNase I (10 U/mL) for 4 h. NET formation was assessed using microscopy or ELISA, as described in the Supplementary Methods.

### NET extraction and cell culture

Mouse bone marrow-derived neutrophils were treated with 80 ng/mL IL-19 for 4 h. The primary renal neutrophils of mice subjected to IP injections of AAI and DNase I were extracted and cultured for an additional 1 h. After the cell culture medium was gently aspirated, NETs adhering to the bottom were washed with 2–3 mL cold PBS and centrifuged at 1000 × *g* for 10 min at 4 °C. The cell-free supernatant containing NETs was gently pipetted and stored at −20 °C after determining the concentration using PicoGreen.

Mouse renal tubular epithelial cells (mRTECs), provided by Prof. Huiyao Lan, were cultured in DMEM/F-12 supplemented with 5% (v/v) heat-inactivated fetal bovine serum at 37 °C in a humidified incubator under 5% CO_2_. The mRTECs were treated with AAI (40 μM) and extracted NETs (500 ng/mL) for 20 h, which were derived from IL-19-treated mouse bone marrow-derived neutrophils or primary renal neutrophils of mice subjected to IP injection of AAI and DNase I.

### Quantification of NETs

To quantify serum NET levels, we used mouse MPO ELISA kits to detect serum MPO levels. Serum nucleosome quantification was performed using the Cell Death kit. DNA levels in free serum and cell supernatants were quantified using the PicoGreen assay kit according to the manufacturer’s instructions.

### Flow cytometry

To assess the level of mRTEC apoptosis, we used an AV-FITC/PI apoptosis detection kit to detect programmed cell death using flow cytometry (Cytoflex, Beckman Coulter, CA, USA), as previously described (49) (Supplementary Methods).

### Acridine orange–ethidium bromide staining

The morphology of apoptotic treated cells was detected and distinguished by acridine orange–ethidium bromide fluorescent staining. Staining was performed as previously reported (8).

### Statistical analysis

Data are expressed as the mean ± standard error of the mean (SEM) from at least three independent experiments. Differences between two groups were compared using a two-tailed Student’s *t*-test. Differences between multiple groups were compared using a one-way analysis of variance (ANOVA) followed by Tukey’s post hoc test. Differences were considered statistically significant at P < 0.05. All analyses were performed using GraphPad Prism (version 8.0; GraphPad Software, San Diego, CA, USA). For further details regarding the materials used, please refer to the Supplementary information.

### Study approval

The in vivo experiments were approved by the Anhui Medical University’s Institutional Animal Care (approval No. LLSC20200730) and Use Committee and performed according to institutional animal care guidelines and conducted in Association for Assessment and Accreditation of Laboratory Animal Care–accredited facilities.

## Acknowledgements

The authors thank the Center for Scientific Research of Anhui Medical University for valuable help in our experiment. Graphical abstract and supplement figure 11 were created by Figdraw.

## Contributions

C.-L.D. performed the cell experiments, analyzed the data, and wrote the manuscript. T.-T.M and C.H. designed, supervised, and wrote the manuscript. J.-H.D. and R.F. provided a series of experimental instructions. Q.W. L.-F.J. and S.-Q.F. performed the animal experiments. C.-T.X. P.-C.J. and Z.-M.Z. assisted with experiments. J.L. designed the experiments. All the authors approved the final version of the manuscript.

## Ethical Statement

All animal experiments were reviewed and approved by the Institutional Animal Experimental Ethics Committee from the Experimental Animal Center of the Anhui Medical University (Hefei, Anhui Province, China). Human investigations were performed in accordance with the Declaration of Helsinki, and approved by the Research Ethics Committee of Anhui Medical University after informed consent was obtained from the patients.

## Data Availability

The datasets are available from the corresponding author on reasonable request.

## Supplementary information for

### Supplementary Materials and Methods

#### Reagents and materials

The sources and catalog numbers of the antibodies used were as follows: kidney injury molecule-1 (KIM-1, bs-2713R), PSTPIP2 (bs-19580R), and β-actin (bs-0061R) from Bioss Antibodies (Beijing, China); citrullinated histone H3 (Cit-H3, ab5103), Ly6G (ab238132), interleukin-19 (IL-19, ab154187), and cleaved caspase-3 (ab214430) from Abcam (Cambridge, UK); PSTPIP2 (13450-1-AP) and E-cadherin (60335-1-Ig) from Proteintech (Wuhan, China); myeloperoxidase (MPO, AF3667) from R&D Systems (Minneapolis, MN, USA); IκB-α (T55026), phospho-IκB-α (Ser32/Ser36, TP56280), NF-κB p65 (T55034), and phospho-NF-κB p65 (Ser536, TP56372) from Abmart (Shanghai, China); and IL-20Rβ (#DF3603) and cleaved caspase-3 (#AF7022) from Affinity Biosciences (Jiangsu, China). Ultra-LEAF™ purified anti-mouse Ly6G Antibody (#127649), Ultra-LEAF™ Purified Rat IgG2a, κ isotype control antibody (#400565), PE anti-mouse/human CD11b antibody (#101208), and APC anti-mouse CD45 antibody (#103112) were purchased from BioLegend (San Diego, CA, USA). Aristolochic acid I (AAI, HY-N0510), GSK484 (HY-100514), and recombinant IL-19 (rIL-19, HY-P7213) were purchased from MedChemExpress (Shanghai, China). DNase I (EN0521) was purchased from Fermentas (Vilnius, Lithuania). Creatinine (Cr) and blood urea nitrogen (BUN) assay kits were purchased from Njjcbio (Nanjing, China). Enzyme-linked immunosorbent assay (ELISA) kits for MPO, MCP-1, tumor necrosis factor (TNF)-α, and IL-19 were purchased from Jymbio (Wuhan, China). An Annexin V-FITC/PI Apoptosis Detection Kit was purchased from BestBio (Shanghai, China). RPMI-1640, Dulbecco’s Modified Eagle’s Medium/Nutrient Mixture F-12 (DMEM/F-12), and heat-inactivated fetal bovine serum were purchased from Gibco (Carlsbad, CA, USA). Mouse bone marrow neutrophil isolation and caspase-3 activity kits were purchased from Solarbio (Beijing, China). A NETosis Assay Kit was purchased from Cayman Chemical (Ann Arbor, MI, USA). A TUNEL Apoptosis Assay Kit and Nuclear and Cytoplasmic Protein Extraction Kit were purchased from Beyotime Biotechnology (Jiangsu, China). A PicoGreen Assay Kit was purchased from Invitrogen (Carlsbad, CA, USA). A Cell Death kit was purchased from Roche (Mannheim, Germany). An enhanced chemiluminescence (ECL) kit was purchased from Thermo Fisher Scientific (Waltham, MA, USA). A Multi Tissue Dissociation Kit 2 and Anti-Ly6G MicroBeads were purchased from Miltenyi Biotec (Bergisch Gladbach, Germany). D-Luciferin potassium salt and a chemiluminescent luciferase substrate were purchased from Abcam (Cambridge, UK).

#### Neutrophil isolation

Mouse bone marrow-derived neutrophils were extracted as follows: Femurs and tibias were aseptically extracted after the animals were sacrificed, and the red marrow cavities were exposed after removing the cartilage at both ends. A 1-mL sterile syringe was used to aspirate a small volume of dilution containing 10% standard fetal bovine serum or serum-containing medium, the marrow cavity was rinsed to obtain marrow, and a single-cell suspension was prepared. Mature neutrophil extraction and purification were performed using the mouse bone marrow neutrophil isolation kit, according to the manufacturer’s instructions.

To isolate and purify mouse renal tissue-derived primary neutrophils, a single-cell suspension was prepared using a gentleMACS™ Dissociator and the Multi Tissue Dissociation Kit 2 (Miltenyi Biotec, Cologne, Germany). We separated and purified the neutrophils using Anti-Ly6G MicroBeads (Miltenyi Biotec) magnetic separation with an autoMACS Pro Separator (Miltenyi Biotec), according to the manufacturer’s instructions. Neutrophils were extracted and cultured for an additional 1 h.

#### Generation of *PSTPIP2* conditional knock-in mice and treatment

To generate kidney-specific *PSTPIP2* knock-in mice, the “CAG promoter-loxP-stop-loxP-mouse *PSTPIP2* CDS-polyA” cassette was inserted into the intron of ROSA26. Expression of mouse *PSTPIP2* CDS is dependent on Cre recombination. The gRNA to ROSA26 gene, the donor vector containing the “CAG promoter-loxP-stop-loxP-mouse *PSTPIP2* CDS-polyA” cassette, and Cas9 mRNA were co-injected into fertilized mouse eggs to generate targeted conditional knock-in offspring. F0 founder animals were identified by PCR followed by sequence analysis and were bred with wild-type mice to test germline transmission and generate F1 animals. All mice were genotyped by PCR before the experiments. *PSTPIP2*^Flox/Flox^ (FF) mouse lines were constructed by Cyagen Biosciences Inc. (Guangzhou, China). *PSTPIP2*-cKI mice were generated by mating *PSTPIP2* FF mice with kidney-specific promoter (Cadherin-16)-driven Cre (KspCre) mice. Only age matched male littermates (8–10 weeks old) were used for the experiments; all mice were genotyped by PCR before experimentation. To generate the AAI-induced AAN model, mice were intraperitoneally injected with a single dose of AAI at 20 mg/kg. We sacrificed 6–10 animals per group under anesthesia 3 d after injection. Kidney tissues and blood were collected for further analysis. Blood was collected for BUN and Cr measurements according to the manufacturer’s instructions. Kidneys were harvested for paraffin embedding, molecular analysis, and immunostaining.

#### AAV9-mediated *PSTPIP2* silencing in *PSTPIP2*-cKI mice

Luciferase-labelled specific kidney tissue locations of AAV9-sh*PSTPIP2* and the corresponding vector were designed by and obtained from Hanheng (Shanghai, China). All *PSTPIP2*-cKI mice were randomly divided into two groups (n = 6 per group): AAV9-NC+AAI and AAV9-sh*PSTPIP2*+AAI. Six *PSTPIP2*^Flox/Flox^ (FF) mice were randomly selected as the AAI injection group. After 3 weeks, a mouse AAN model was established 3 d after AAV9 administration. Mice exposed to AAV9 were anesthetized, and the effect of AAV9-sh*PSTPIP2* on kidney tissue localization was confirmed using an IVIS Lumina III Imaging System (Caliper Life Sciences, Hopkinton, MA, USA). AAV9-NC+AAI mice were injected with an empty rAAV9 vector via the tail vein, and AAI was administered at 10 mg/kg in a single intraperitoneal injection. AAV9-sh*PSTPIP2*+AAI mice were injected with an AAV9-sh*PSTPIP2* vector via the tail vein, and AAI was administered at 10 mg/kg in a single intraperitoneal injection. Mice were sacrificed 72 h after AAI injection, and blood and kidney tissues were collected.

#### Caspase-3 activity assay

The activity of caspase-3 was measured using caspase-3 activity kits. Assays were performed in 96-well microtiter plates, to which 50 μL protein extract, 40 μL reaction buffer, and 10 μL caspase substrate were added sequentially. The protein extracts were incubated at 37 °C for 1–2 h. The absorbance of the samples was measured using a BioTek Cytation 5 Cell Imaging Multimode Reader (BioTek, Winooski, VT, USA) at 405 nm.

#### Serum Cr and BUN assays

The serum concentrations of Cr and BUN were determined using the corresponding assay kits, according to the manufacturer’s instructions.

#### Flow cytometry

Harvested cells were stained with 400 μL Annexin V binding solution and resuspended. Then, 5 μL annexin V-FITC solution was added, the solution was kept in the dark at 4 °C for 15 min, 10 μL propidium iodide was added, and the solution was left for an additional 5 min in the dark at 4 °C after gentle mixing. The solution was examined with a flow cytometer and analyzed using CytExpert 2.1 (Beckman Coulter). Neutrophil infiltration (CD11b^+^Ly6G^+^) was analyzed by flow cytometry as previously described (20).

#### TUNEL assay

Renal cell apoptosis was examined by the TUNEL assay using a One-step TUNEL Apoptosis Assay Kit (Beyotime Biotechnology). Briefly, cells were fixed with 4% paraformaldehyde in PBS and exposed to a TUNEL reaction mixture containing TM green–labeled dUTP. The samples were counterstained with 4′,6-diamidino-2-phenylindole (DAPI, Servicebio, Wuhan, China). TUNEL^+^ nuclei were identified using fluorescence microscopy (Olympus, Tokyo, Japan).

#### Histopathology

Mouse renal tissues were fixed in 4% paraformaldehyde for 24 h immediately after excision, processed for histological examination according to a conventional method, and stained with hematoxylin and eosin (H&E; Servicebio). Hematoxylin-stained tissues were visualized and photographed using an automated digital slide scanner (Pannoramic MIDI, 3DHISTECH, Budapest, Hungary).

#### ELISA

After blood collection, the serum and erythrocytes were rapidly and carefully separated by centrifugation at 3000 × *g* for 10 min. The tissues were mashed with the addition of an appropriate amount of saline. The supernatant was removed by centrifugation at 3000 rpm for 10 min. Serum and tissue monocyte chemoattractant protein-1 (MCP-1) and TNF-α levels were measured using ELISA kits according to the manufacturer’s instructions. mRTECs were treated either with AAI (40 µM) or saline for 20 h to evaluate the concentration of IL-19 in the supernatant via ELISA. The concentration of IL-19 in the sera of AAN mice was also measured by ELISA using a commercial protocol.

#### NET formation microscopic analysis

For microscopic analysis, treated bone marrow-derived neutrophils or renal tissue-derived primary neutrophils were plated on poly L-lysine-coated cover slips. Neutrophils were incubated overnight at 4 °C with antibodies against Cit-H3 (1:100) followed by goat anti-rabbit IgG for 1 h at room temperature (15–25 °C). The cells were counterstained with DAPI and visualized using fluorescence microscopy. The levels of neutrophil elastase, which forms part of NETs, were measured using the Cayman Chemical NETosis assay kit according to the manufacturer’s specifications. The levels of dsDNA, which forms another part of NETs, were measured using PicoGreen, according to the manufacturer’s specifications.

#### Immunohistochemistry

Kidney tissues were fixed in 4% paraformaldehyde for 24 h immediately after excision, embedded in paraffin, and cut to 5-μm-thick sections. The sections were exposed to 3% hydrogen peroxide for 10 min to suppress endogenous peroxidase activity and then blocked with 3% bovine serum albumin (BSA; Servicebio) for 30 min. The tissue sections were incubated with anti-Ly6G (1: 2,000), KIM-1 (1:200), and cleaved caspase-3 (1:100) antibodies overnight at 4 °C. The sections were then washed and incubated with anti-rabbit IgG secondary antibodies for 30 min at room temperature (15–25 °C). The sections were counterstained with hematoxylin and developed using a diaminobenzidine (DAB) kit. The stained sections were photographed and visualized using the Pannoramic MIDI (3DHISTECH).

#### Immunofluorescence staining

Mouse renal tissue sections were blocked with 5% BSA at 37 °C for 30 min to avoid non-specific staining. Sections were incubated with anti-Cit-H3 (1:100), anti-MPO (1:100), anti-IL-19 (1:100), anti-E-cadherin (1:100), or anti-PSTPIP2 (1:100) antibodies overnight at 4 °C. Then, the sections were incubated with a secondary antibody (1:100) in the dark at 37 °C for 1 h. After the nuclei were stained with DAPI, the stained sections were examined by inverted fluorescence microscopy (Olympus).

After discarding the cell culture medium, mRTECs were washed twice with PBS, blocked with 5% BSA for 30 min after fixation and permeabilization, and incubated with anti-KIM-1 (1:100), PSTPIP2 (1:100), IL-19 (1:100), and NF-κB p65 (1:50) antibodies at 4 °C overnight. Neutrophils were incubated overnight at 4 °C with antibodies against Cit-H3 (1:100) and IL-20Rβ (1:200) antibodies. After washing with PBS, cells were incubated with the corresponding fluorescent secondary antibodies for 1 h at room temperature (15–25 °C). After the nuclei were stained with DAPI (Servicebio), the sections were blocked with an anti-fluorescent quencher-containing blocking solution and observed under a fluorescence microscope (Olympus).

#### Western blot analysis

Cells were lysed with radioimmunoprecipitation assay (RIPA) lysis buffer to extract proteins from mouse kidney tissues (20 or 40 mg), mRTECs, and neutrophils. The Nuclear and Cytoplasmic Protein Extraction Kit (Beyotime Biotechnology) was used to extract nuclear and cytoplasmic proteins from 6 × 10^5^ mRTECs, according to the manufacturer’s instructions. Whole extracts were separated using 8–12% sodium dodecyl sulfate polyacrylamide gel electrophoresis (SDS-PAGE), transferred to a polyvinylidene fluoride (PVDF) membrane (Millipore, Billerica, MA, USA), blocked with 5% non-fat dry milk (NFDM) for 2 h at room temperature (15–25 °C), and washed thrice with Tris-buffered saline solution. PVDF membranes were incubated with primary antibodies against KIM-1, cleaved caspase-3, Cit-H3, PSTPIP2, IL-20Rβ, NF-κB p65, phospho-NF-κB p65 (Ser536), IκB-α, phospho-IκB-α (Ser32/Ser36), Histone H3, and β-actin. The membranes were washed with TBS-Tween 20 and incubated with secondary antibodies. After extensive washing in TBS-Tween 20, protein bands were visualized with the ECL kit (Epizyme, Shanghai, China). The signal intensity of each protein band was quantified using ImageJ software (National Institutes of Health, Bethesda, MD, USA).

#### Real-time reverse transcription-PCR

Total RNA was extracted from kidney tissues using TRIzol reagent (Invitrogen). First-strand cDNA was synthesized using a ThermoScript RT-PCR synthesis kit (AG, Hunan, China), according to the manufacturer’s instructions. Real-time quantitative PCR analysis of mRNA was performed using ThermoScript RT-qPCR kits in an ABI Prizm step-one plus real-time PCR system (Applied Biosystems, Foster City, CA, USA). The products were used as templates for amplification using SYBR Green PCR amplification reagent and gene-specific primers. The relative expression levels were calculated using the standard 2^−ΔΔCt^ method. The forward and reverse primers used for PCR are listed in Table S1.

#### Clinical specimens

The clinical features of patients with various kidney diseases are listed in table S2. This study was approved by the Biomedical Ethics Committee of the Anhui Medical University (ethical clearance no.: 20200734). The experiments were conducted with the understanding and written consent of each patient.

#### Transwell migration assay of neutrophil levels

Isolated bone marrow-derived neutrophils were resuspended in serum-free RPMI-1640 medium containing 1% penicillin/streptomycin (P/S) at 10^6^/mL. Then, 200 μL of cells were seeded in the upper well of a Transwell chamber with microporous filters (3-μm pores; Corning, Corning, NY, USA), and 500 μL of 1% P/S serum-free RPMI-1640 medium in the absence or presence of IL-19 (80 nM) was added to the bottom chamber. After incubation for 3 h, cells adherent to the membrane in a 3-μm pore Transwell system were fixed with 4% paraformaldehyde for 10 min, stained with 1% crystal violet for 30 min, and observed under a light microscope (Olympus).

#### Transfection with *PSTPIP2* plasmid

mRTECs (2–4 × 10^5^ cells) were seeded into a 6-well plate and cultured in DMEM/F-12 medium (containing 5% FBS) until cell attachment and grown to a density of ∼50%. Cells were transfected with 2000 ng/mL pEX-2-Control or pEX-2-*PSTPIP2* overexpression plasmid mixed with a Lipo3000 transfection reagent (Hanheng). After 6 h, the Opti-MEM was replaced with DMEM/F-12 containing 5% FBS, and the cells were treated with AAI (40 μM) for 20 h. Transfection efficiency was determined using western blotting and real-time PCR.

#### Co-immunoprecipitation

Cells were collected after washing with pre-chilled PBS solution and an appropriate amount of pre-chilled lysis buffer was added. After ultrasonic disruption and centrifugation, the cell supernatants were collected and stored. *PSTPIP2* and NF-κB p65 antibodies (1–5 ug) were added to the samples, along with a nonspecifically immunized cognate antibody as a control. After overnight incubation at 4 °C, 5 μL of protein A-agarose beads and 5 μL protein G-agarose beads were added to the samples and mixed gently overnight at 4 °C. Immunocomplex pellets were obtained after centrifugation, resuspended in SDS buffer, boiled, stored, and resolved in 8–12% SDS-polyacrylamide gels for western blot analysis with PSTPIP2 and NF-κB p65 antibodies.

**Table S1.**
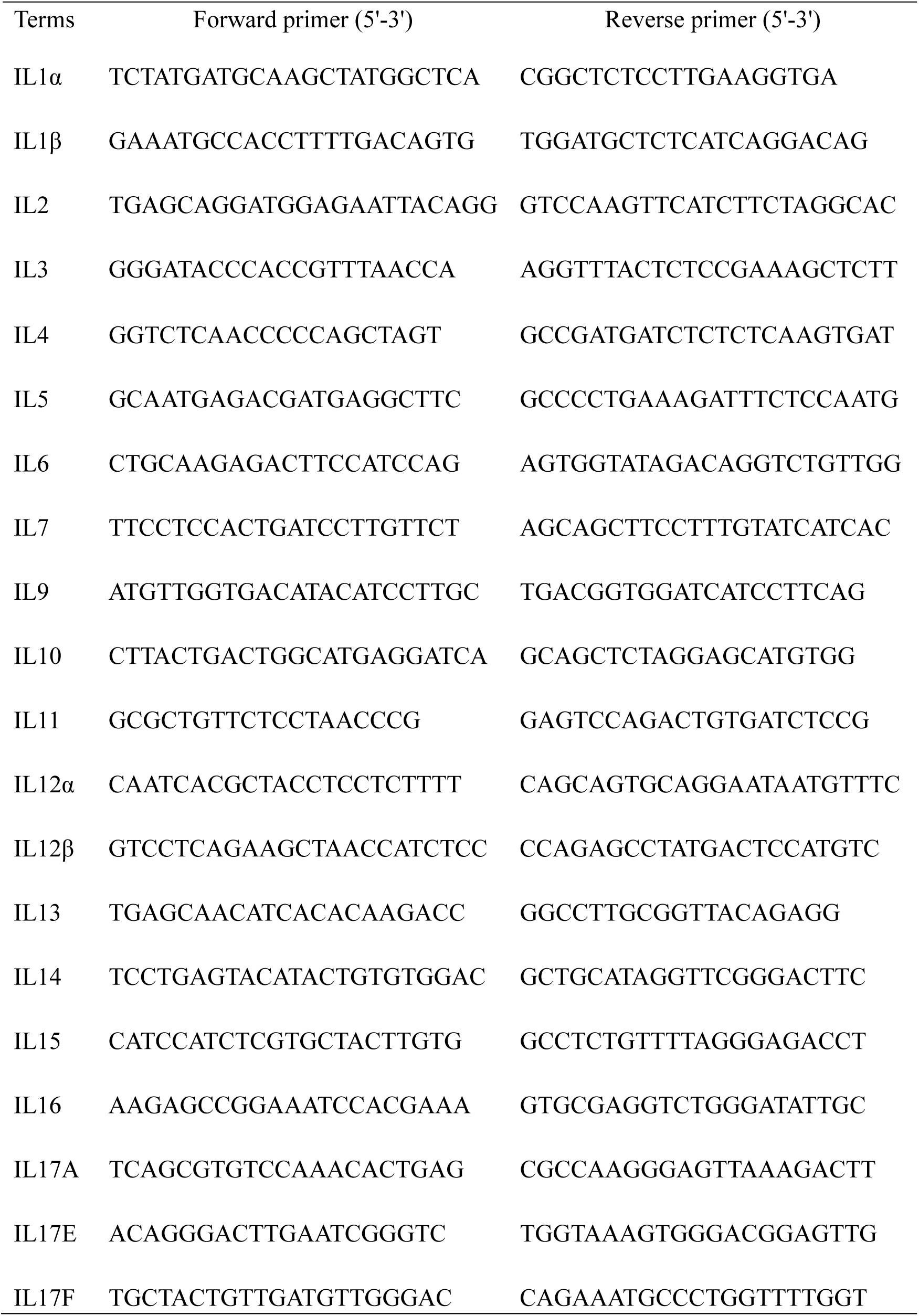
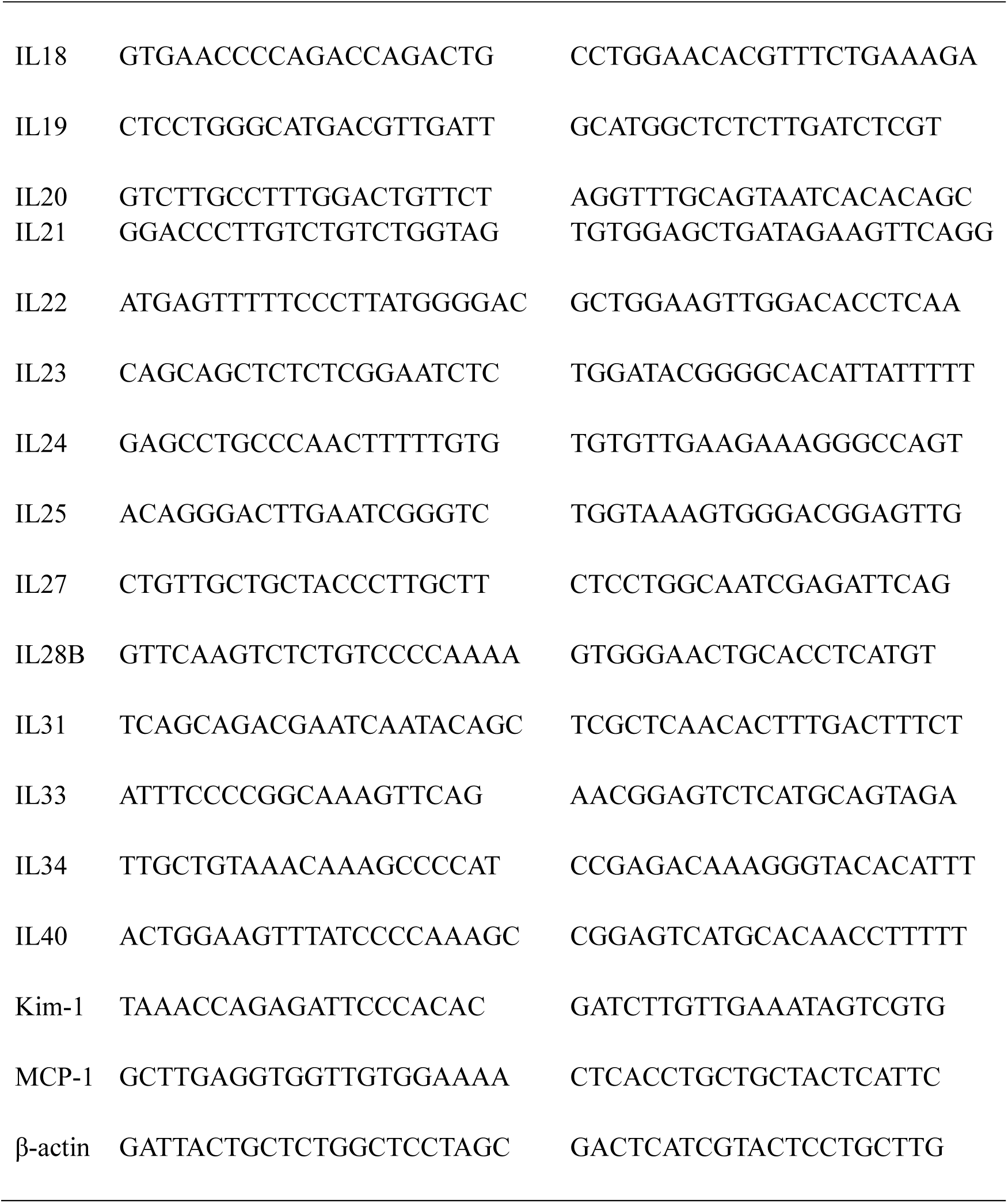
Primer sequences for the quantitative real-time PCR analysis of mouse tissues.

**Table S2.**
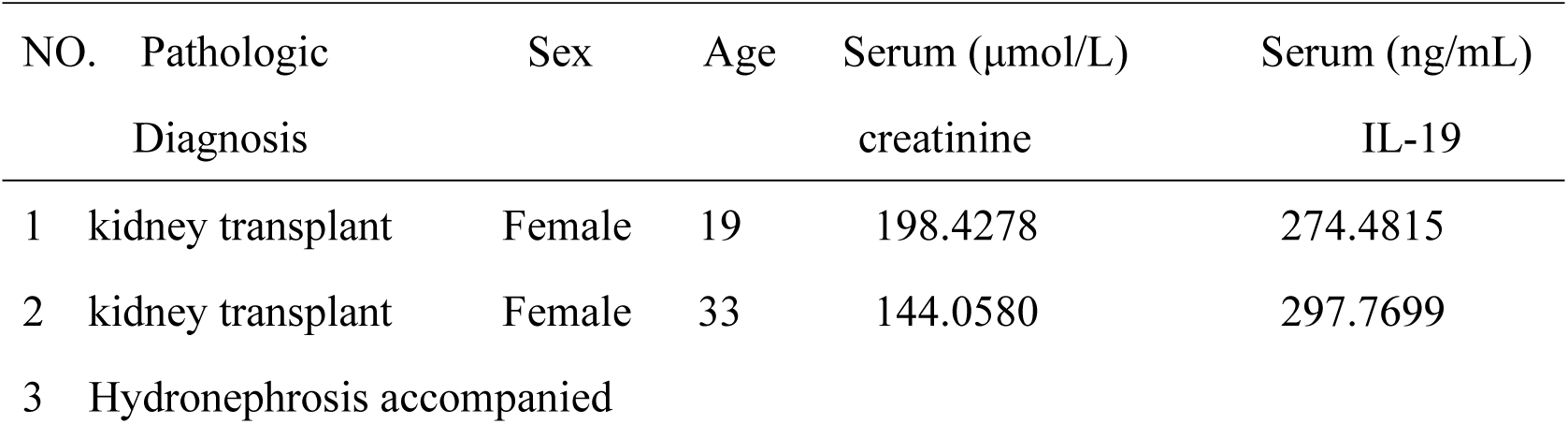
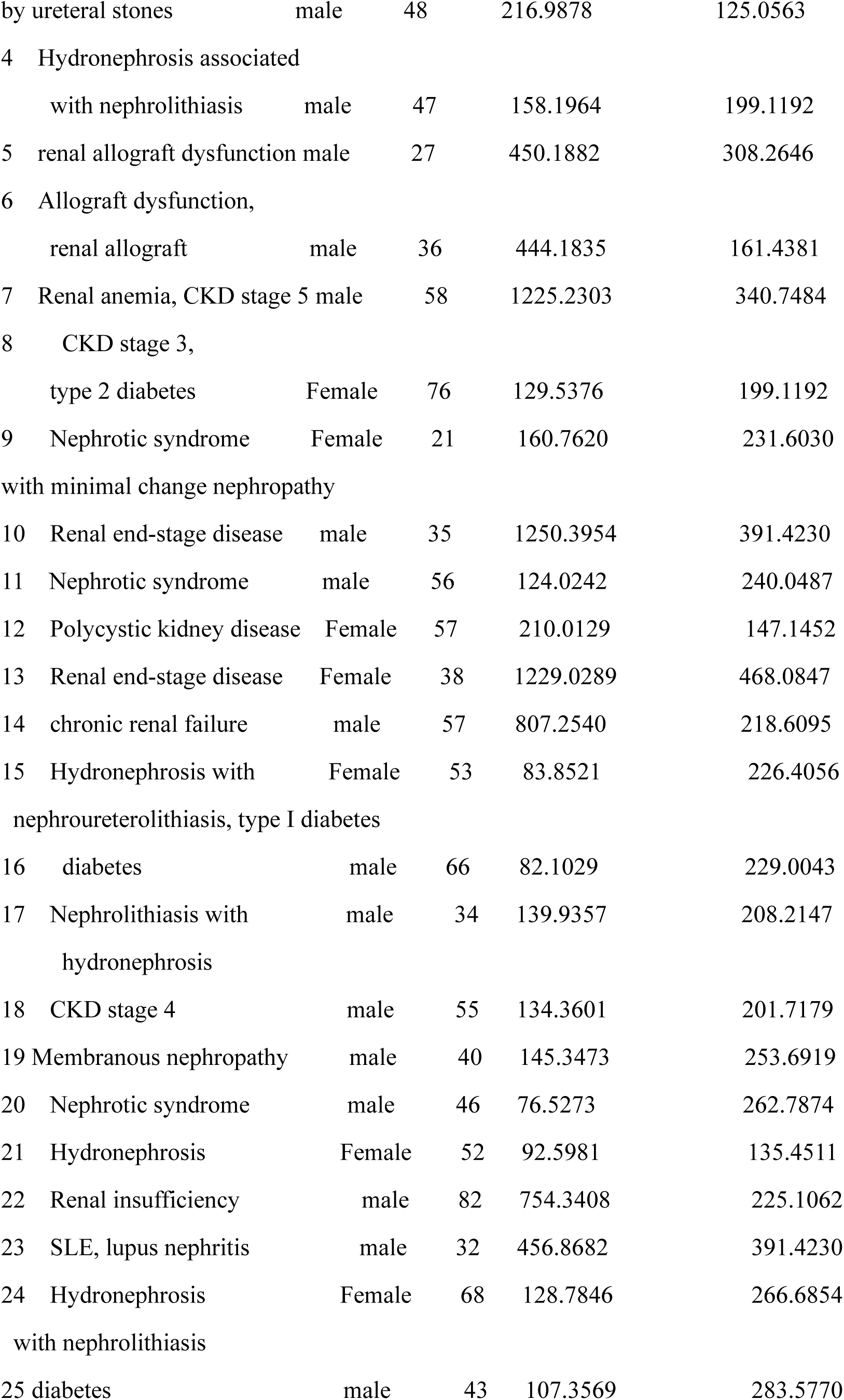
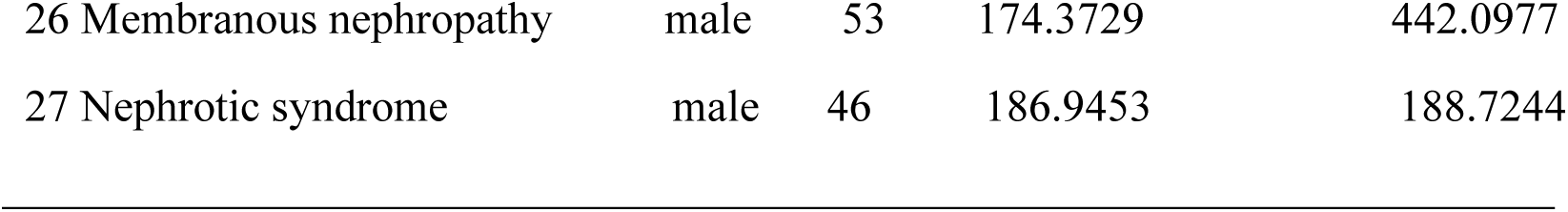
Clinical features of the patients with kidney diseases.

**Figure S1.**
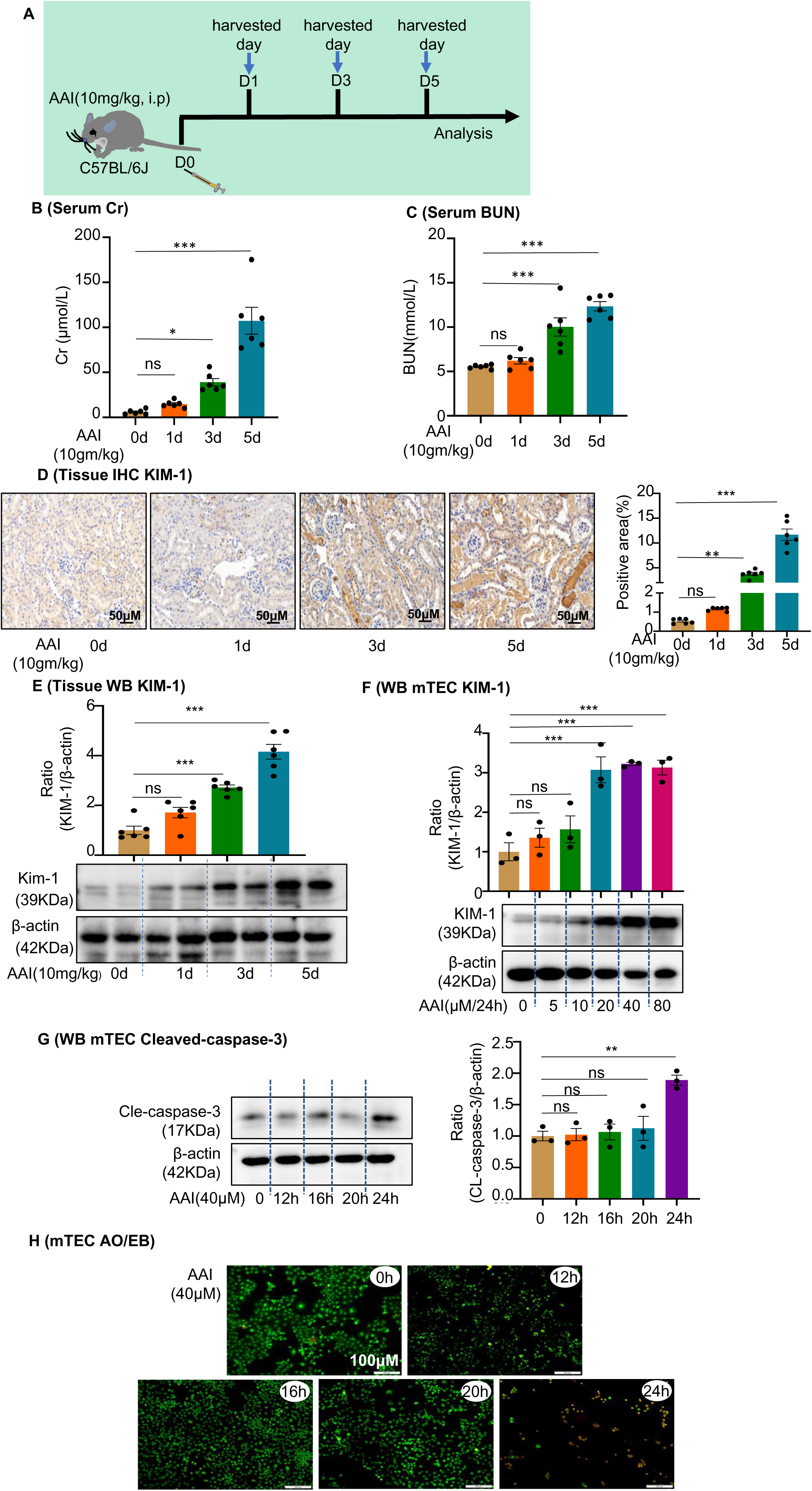
Aristolochic acid I (AAI)-induced acute aristolochic acid nephropathy (AAN) *in vivo* model and mouse renal tubular epithelial cell (mRTEC) injury *in vitro* model. **(A)** Establishment of AAI-induced acute AAN mouse model (n = 6–8 per group). **(B)** Serum creatinine (Cr) assay at various times after AAI treatment (n = 6). **(C)** Blood urea nitrogen (BUN) assay at various times after AAI treatment (n = 6). **(D)** Immunohistochemistry of kidney injury molecule-1 (KIM-1) in different groups of kidney tissues. Positively stained areas were measured using Image-Pro Plus 6.0 (Media Cybernetics, Inc., Rockville, MD, USA) (n = 6). Scale bar, 50 μm. **(E)** Protein level of KIM-1 assessed by western blotting at various times after AAI treatment (n = 6). **(F)** Protein level of KIM-1 after treatment with different AAI concentrations (n = 3). **(G)** Level of cleaved caspase-3 at different times after AAI (40 μM) treatment (n = 3). **(H)** Apoptosis was assessed by acridine orange–ethidium bromide staining and counting of apoptotic nuclei (n = 3). Scale bar, 100 μm. Data are presented as the mean ± SEM of three to six biological replicates per condition. Each dot represents a sample. Statistically significant differences were determined by an independent sample *t*-test and one-way ANOVA followed by Tukey’s post hoc test. *P < 0.05, **P < 0.01, ***P < 0.001, ns: non-significant. **Figure S1—source data 1** **Figure S1—source data 2** **Aristolochic acid I (AAI)-induced acute aristolochic acid nephropathy (AAN) *in vivo* model and mouse renal tubular epithelial cell (mRTEC) injury *in vitro* model.**

**Figure S2.**
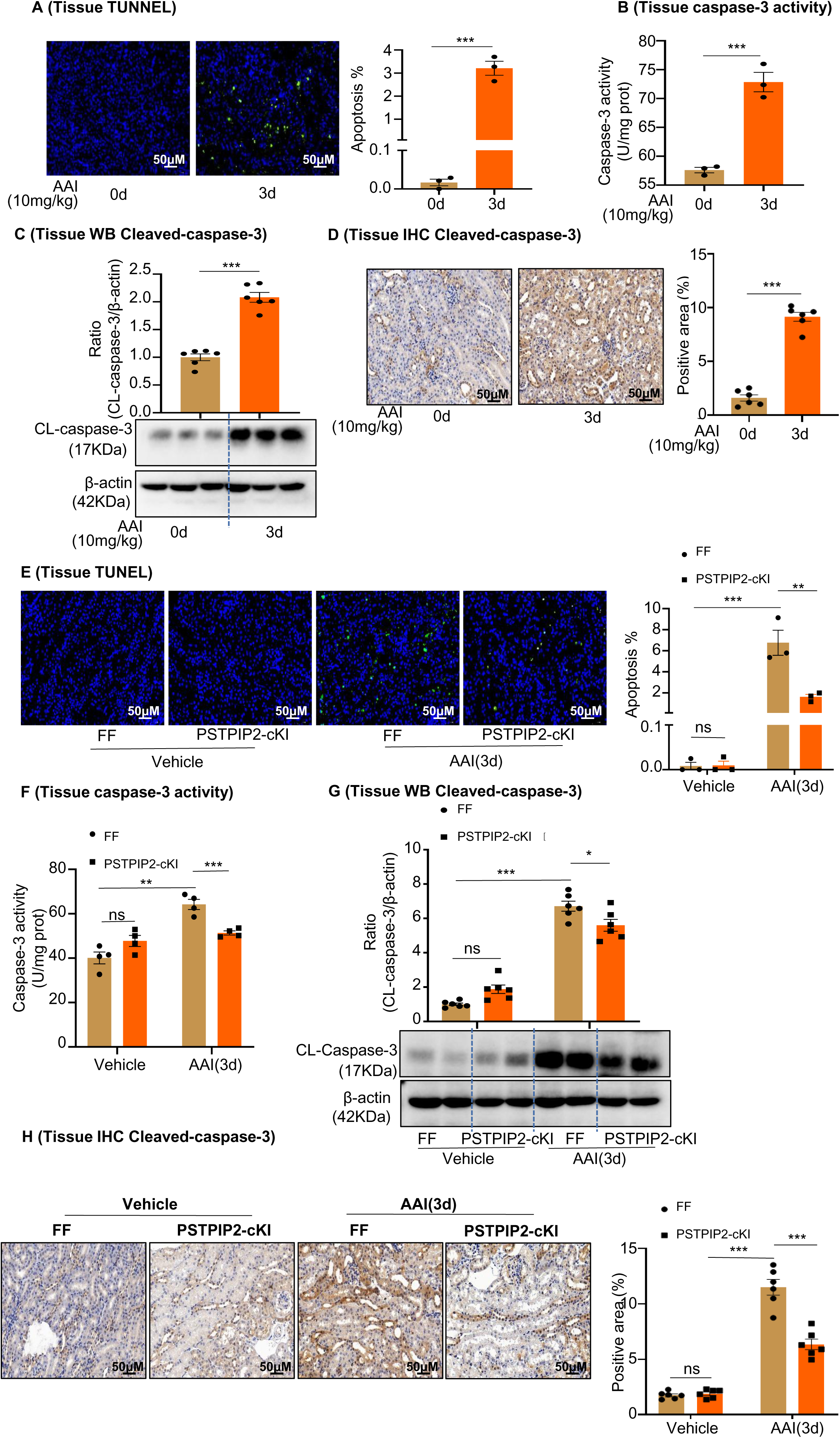
*PSTPIP2*-cKI reduced aristolochic acid I (AAI)-induced kidney apoptosis. **(A)** TUNEL staining of kidney sections after AAI treatment for 3 d and quantification (n = 3). Scale bar, 50 μm. **(B)** Caspase-3 activity in kidney tissues quantified after AAI treatment for 3 d (n = 3). **(C)** Protein levels of cleaved caspase-3 assessed by western blotting after AAI treatment for 3 d (n = 6). **(D)** Immunohistochemical (IHC) analysis and quantification of cleaved caspase-3 in kidney tissues after AAI treatment for 3 d (n = 6). Scale bar, 50 μm. **(E)** TUNEL staining of kidney sections from *PSTPIP2*^Flox/Flox^ (FF) and *PSTPIP2*-cKI mice treated with AAI and quantification (n = 3). Scale bar, 50 μm. **(F)** Quantification of caspase-3 activity in kidney tissues from FF and *PSTPIP2*-cKI mice treated with AAI (n = 4). **(G)** Protein level of cleaved caspase-3 assessed by western blotting (n = 6). **(H)** IHC analysis and quantification of cleaved caspase-3 in kidney tissues from FF and *PSTPIP2*-cKI mice treated with AAI (n = 6). Scale bar, 50 μm. Data are presented as the mean ± SEM of three to six biological replicates per condition. Each dot represents a sample. Statistically significant differences were determined by an independent sample *t*-test and one-way ANOVA followed by Tukey’s post hoc test. *P < 0.05, **P < 0.01, ***P < 0.001, ns: non-significant. **Figure S2—source data 1** **Figure S2—source data 2** ***PSTPIP2*-cKI reduced aristolochic acid I (AAI)-induced kidney apoptosis.**

**Figure S3.**
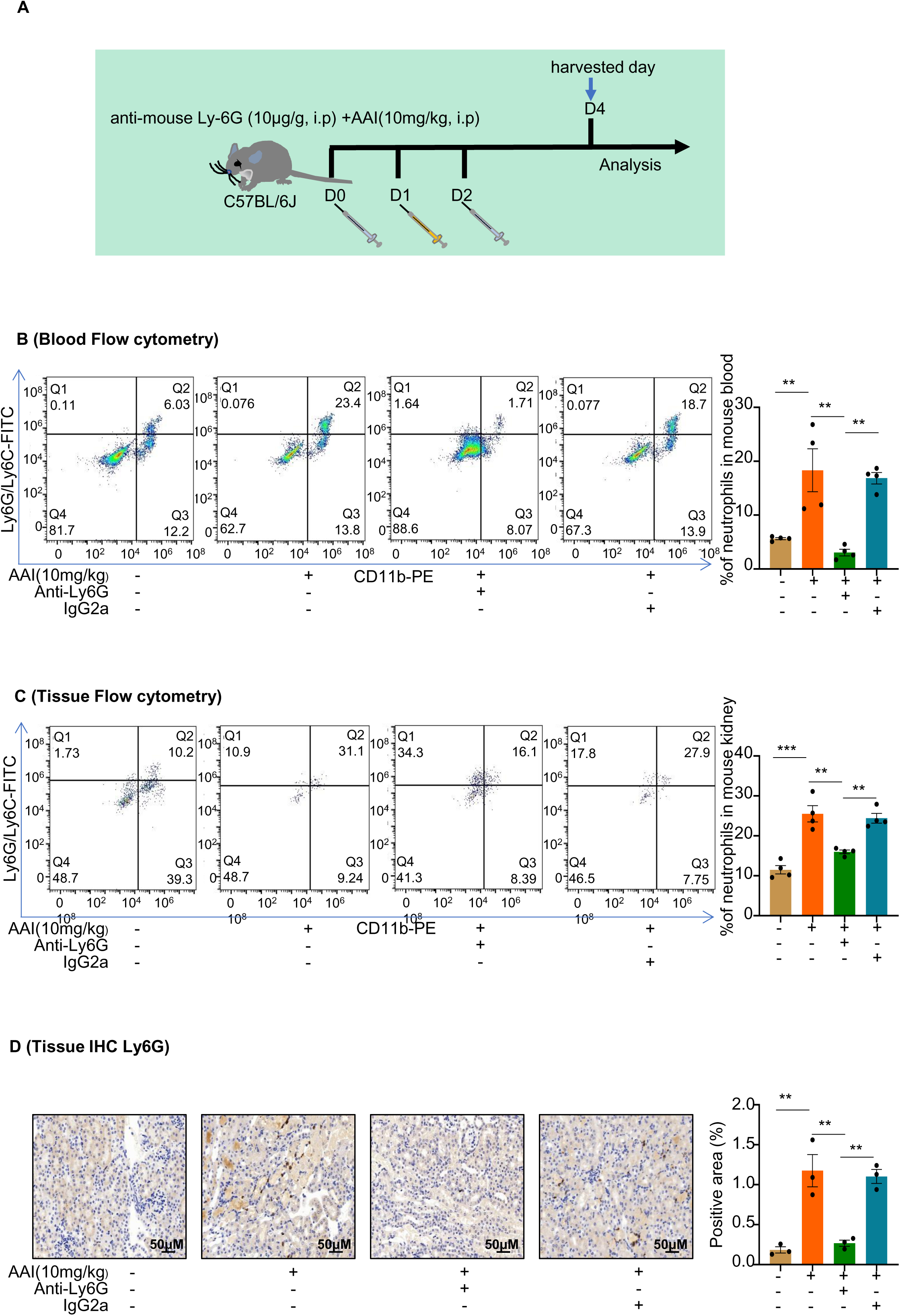
Mice were pretreated with anti-mouse Ly6G or IgG control antibody followed by aristolochic acid I (AAI) treatment. **(A)** Schematic of intraperitoneal (IP) injection of anti-mouse Ly6G neutralizing antibody or IgG control antibody; purified anti-mouse Ly6G (10 µg/g mouse), and IgG control antibody (IP 10 µg/g mouse) was administered daily one day before and after IP AAI injection. **(B and C)** Flow cytometry was used to evaluate Ly6G^+^ neutrophil infiltration in the blood and renal samples. Representative images of flow cytometry analysis are shown in B and C (n = 4 per group). **(D)** Representative images of immunohistochemistry (IHC) staining of Ly6G. Scale bar, 50 μm. Quantification of the percentage of Ly6G^+^ neutrophil infiltration (n = 3 per group). Data are presented as the mean ± SEM of three to six biological replicates per condition. Each dot represents a sample. Statistically significant differences were determined by one-way ANOVA followed by Tukey’s post hoc test. **P < 0.01, ***P < 0.001. **Figure S3—source data 1** **Mice were pretreated with anti-mouse Ly6G or IgG control antibody followed by aristolochic acid I (AAI) treatment.**

**Figure S4.**
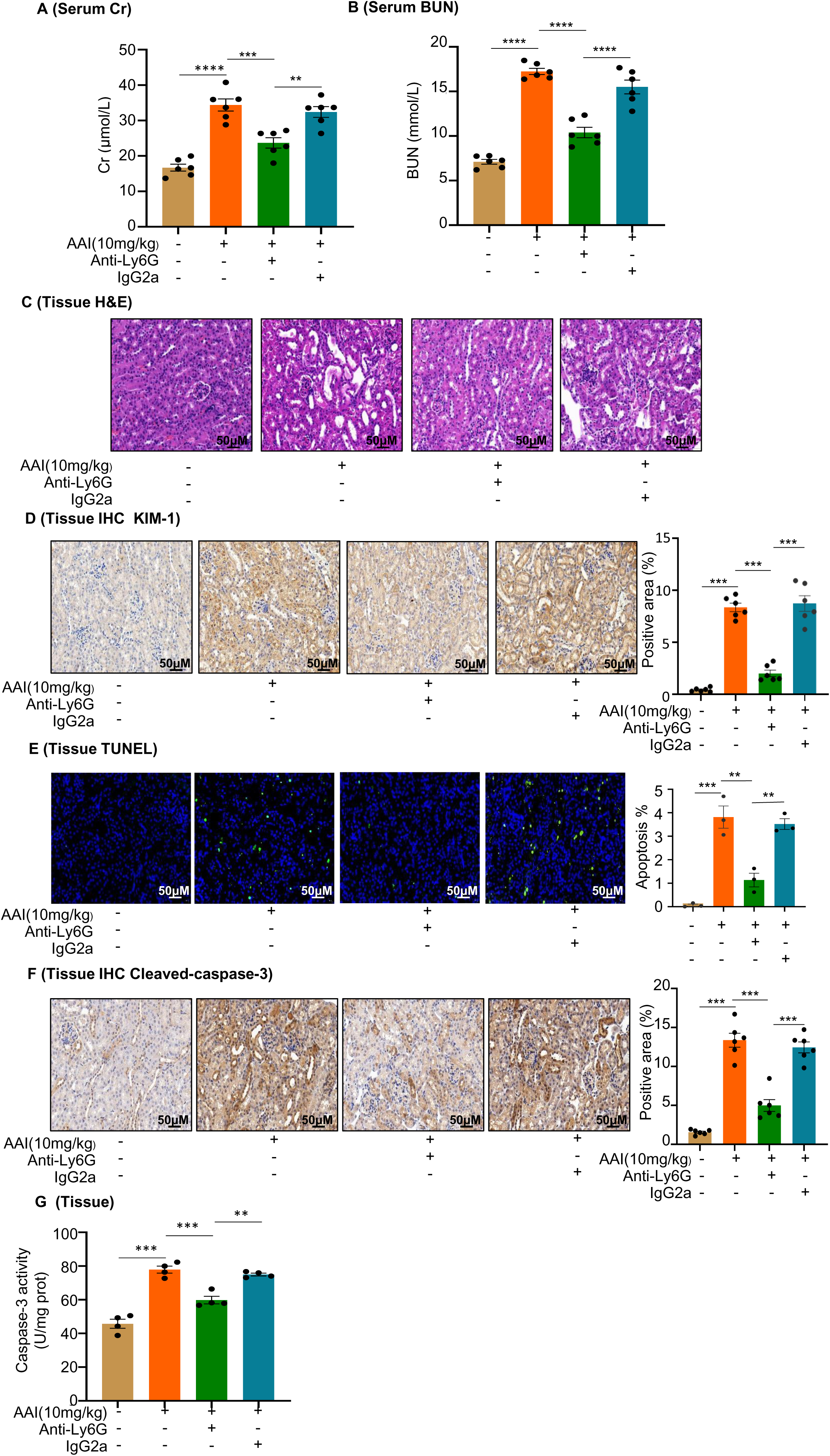
*In vivo* neutrophil depletion attenuates aristolochic acid I (AAI)-induced kidney damage in mice. **(A and B)** Serum creatinine (Cr) and blood urea nitrogen (BUN) assay of mice pretreated with anti-mouse Ly6G or IgG control antibody, followed by AAI treatment (n = 6 per group). **(C)** Representative H&E-staining images of kidney tissues (n = 6 per group). Scale bar, 50 μm. **(D)** Immunohistochemistry (IHC) staining and quantification of kidney injury molecule-1 (KIM-1) in kidney tissues (n = 6 per group). Scale bar, 50 μm. **(E)** TUNEL staining and quantification in kidney sections (n = 3 per group). Scale bar, 50 μm. **(F)** IHC staining and quantification of cleaved caspase-3 in kidney tissues. Scale bar, 50 μm (n = 6 per group). **(G)** Detection of caspase-3 activity in kidney tissues (n = 4 per group). Data are presented as the mean ± SEM of three to six biological replicates per condition. Each dot represents a sample. Statistically significant differences were determined by one-way ANOVA followed by Tukey’s post hoc test. **P < 0.01, ***P < 0.001, ****P < 0.0001. **Figure S4—source data 1** ***In vivo* neutrophil depletion attenuates aristolochic acid I (AAI)-induced kidney damage in mice.**

**Figure S5.**
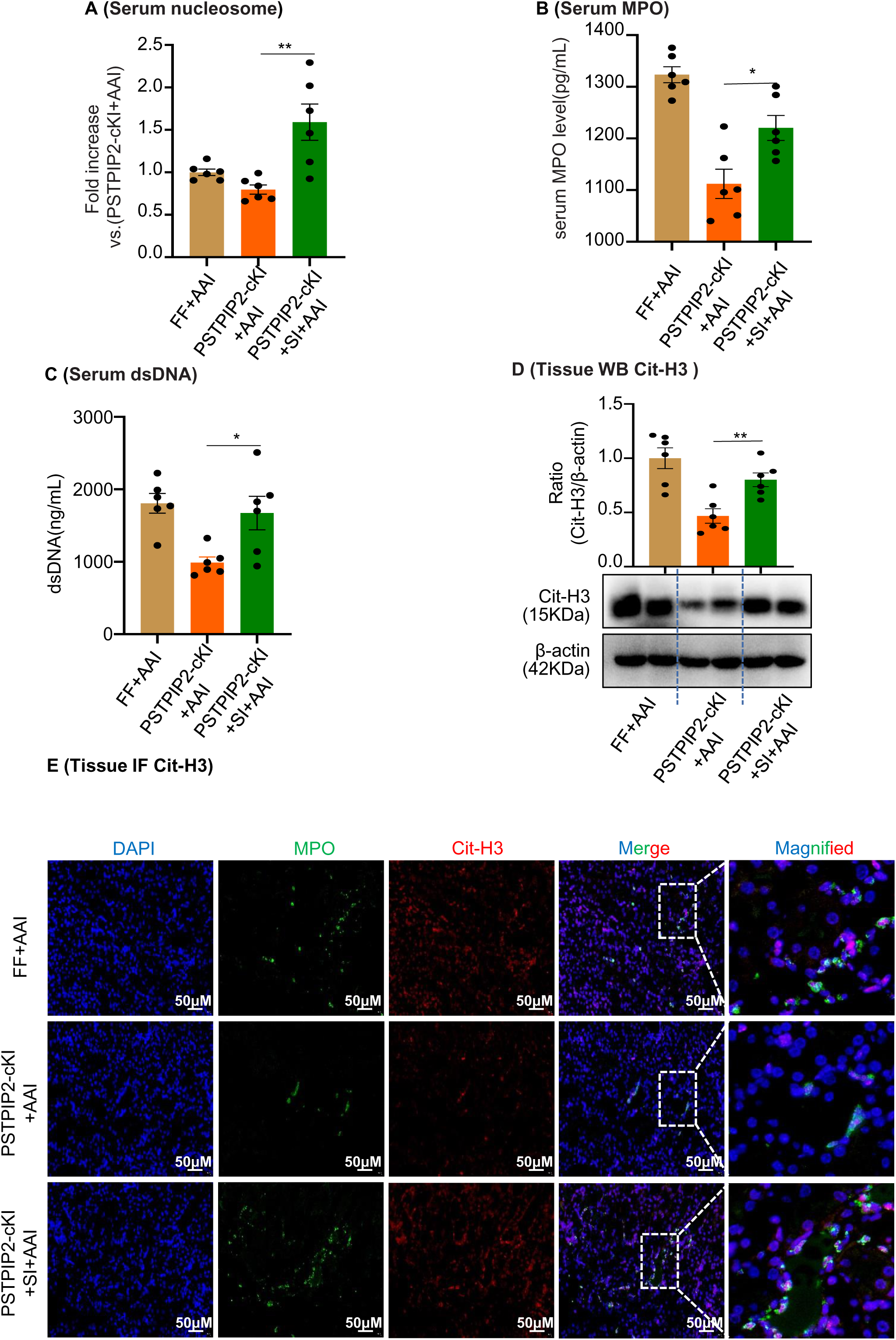
AAV9-mediated *PSTPIP2* restoration re-induces neutrophil extracellular trap (NET) formation in *PSTPIP2*-cKI mice. **(A)** Serum nucleosome level (n = 6 per group). **(B)** Serum myeloperoxidase (MPO) level (n = 6 per group). **(C)** Serum dsDNA level (n = 6 per group). **(D)** Protein level of citrullinated histone 3 (Cit-H3) assessed by western blotting (n = 6 per group). **(E)** Immunofluorescence staining revealed increased colocalization of MPO (green) and Cit-H3 (red, n = 6 per group). Scale bar, 50 μm. Data are presented as the mean ± SEM of three to six biological replicates per condition. Each dot represents a sample. Statistically significant differences were determined by an independent sample *t*-test and one-way ANOVA followed by Tukey’s post hoc test. *P < 0.05, **P < 0.01. **Figure S5—source data 1** **Figure S5—source data 2** **AAV9-mediated *PSTPIP2* restoration re-induces neutrophil extracellular trap (NET) formation in *PSTPIP2*-cKI mice.**

**Figure S6.**
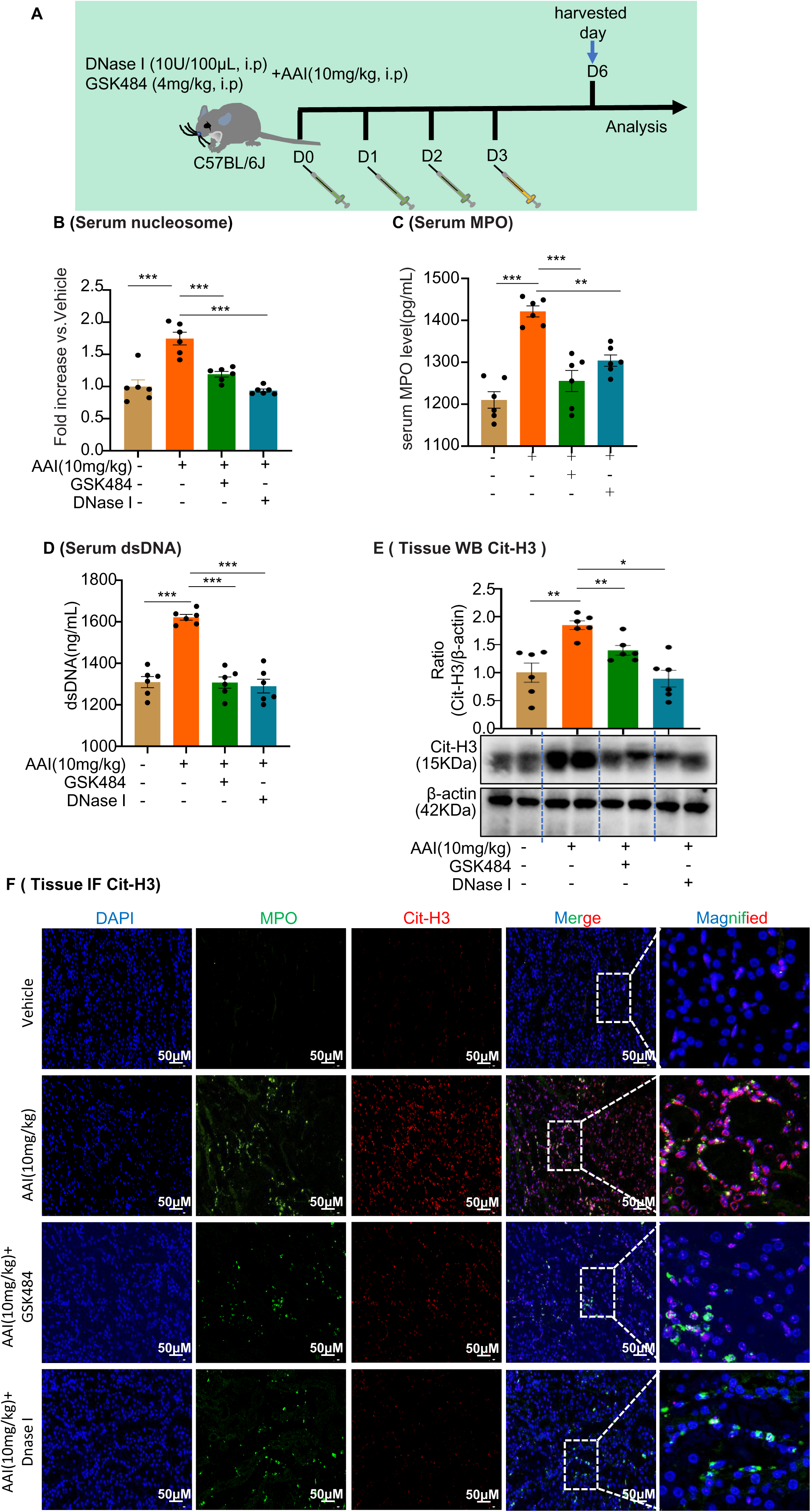
Neutrophil extracellular trap (NET) formation inhibitor, GSK484, or DNase I inhibited formation of NETs in aristolochic acid I (AAI)-treated mice. **(A)** Schematic of GSK484 or DNase I injection in mice experiments. DNase I (10 U/100 μL, 0.9% NaCl) or GSK484 (4 mg/kg) was administered daily for 3 d before AAI treatment. **(B and C)** Serum nucleosome and myeloperoxidase (MPO) level assessed after treatment with AAI and co-treatment with GSK484 or DNase I (n = 6 per group). **(D)** Serum dsDNA level assessed after treatment with AAI and co-treatment with GSK484 or DNase I (n = 6 per group). **(E)** Protein level of citrullinated histone 3 (Cit-H3) assessed by western blotting (n = 6 per group). **(F)** Representative IF staining images of kidney sections, showing the expression of Cit-H3 (red) and MPO (green, n = 6 per group). Scale bar, 50 μm. Data are presented as the mean ± SEM of three to six biological replicates per condition. Each dot represents a sample. Statistically significant differences were determined by an independent sample *t*-test and one-way ANOVA followed by Tukey’s post hoc test. *P < 0.05, **P < 0.01, ***P < 0.001. **Figure S6—source data 1** **Figure S6—source data 2** **Neutrophil extracellular trap (NET) formation inhibitor, GSK484, or DNase I inhibited formation of NETs in aristolochic acid I (AAI)-treated mice.**

**Figure S7.**
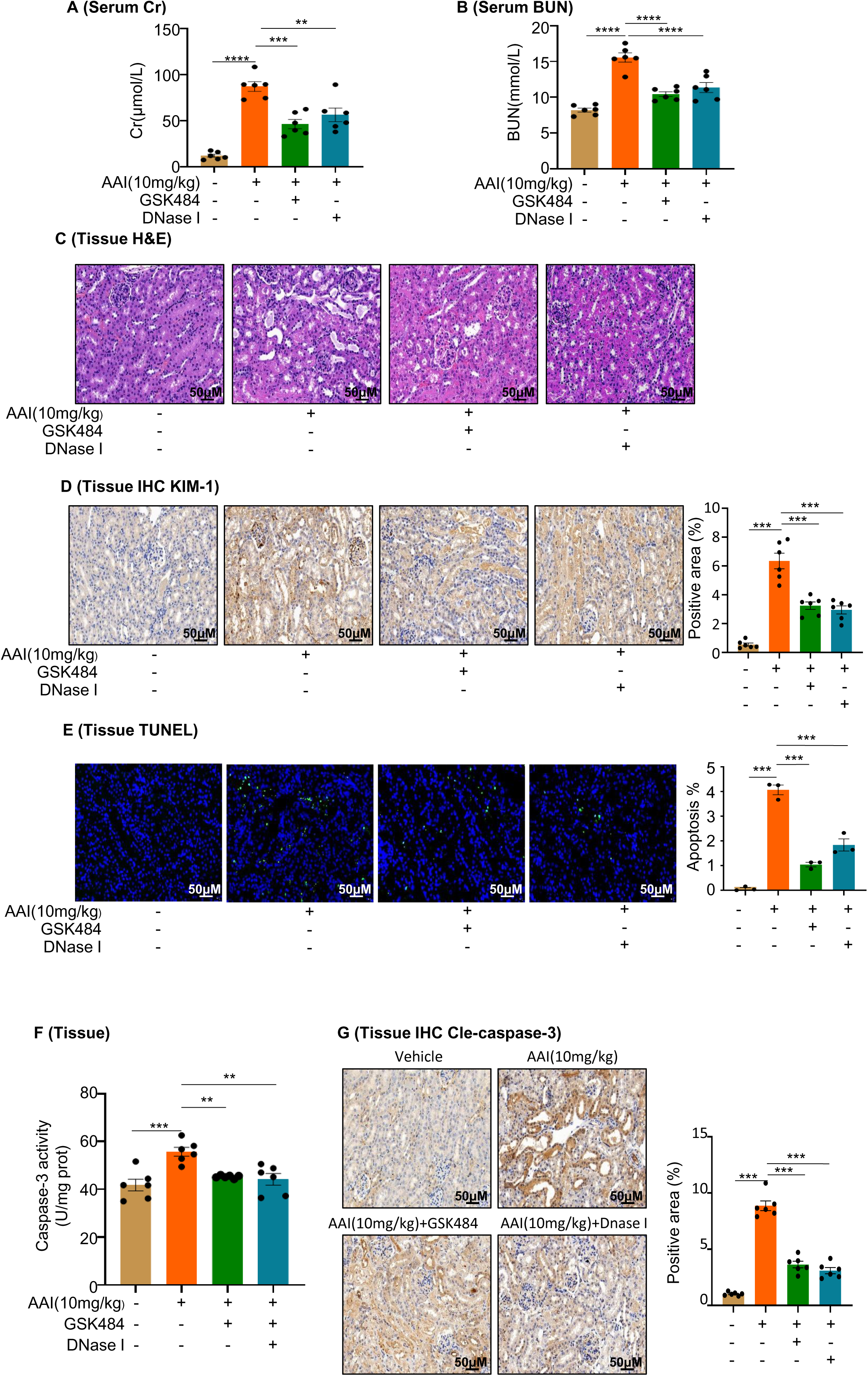
Inhibition of neutrophil extracellular trap (NET) formation attenuates aristolochic acid I (AAI)-induced kidney damage and apoptosis. **(A and B)** Serum creatinine (Cr) and blood urea nitrogen (BUN) assay performed on AAI-treated mice co-treated with GSK484 or DNase I (n = 6 per group). **(C)** H&E staining of kidney sections of AAI-treated mice co-treated with GSK484 or DNase I (n = 6 per group). Scale bar, 50 μm. **(D)** KIM-1 staining of kidney sections of AAI-treated mice co-treated with GSK484 or DNase I (n = 6 per group). Scale bar, 50 μm. Positive staining areas were measured using Image-Pro Plus 6.0 (Media Cybernetics, Inc., Rockville, MD, USA). **(E)** TUNEL staining and quantification of kidney sections (n = 3 per group). Scale bar, 50 μm. **(F)** Activity of caspase-3 quantified after co-treatment with AAI and GSK484 or DNase I (n = 6 per group). **(G)** Cleaved-caspase-3 staining and quantification in kidney sections of AAI-treated mice co-treated with GSK484 or DNase I (n = 6 per group). Scale bar, 50 μm. Data are presented as the mean ± SEM of three to six biological replicates per condition. Each dot represents a sample. Statistically significant differences were determined by one-way ANOVA followed by Tukey’s post hoc test. **P < 0.01, ***P < 0.001, ****P < 0.0001. **Figure S7—source data 1** **Inhibition of neutrophil extracellular trap (NET) formation attenuates aristolochic acid I (AAI)-induced kidney damage and apoptosis.**

**Figure S8.**
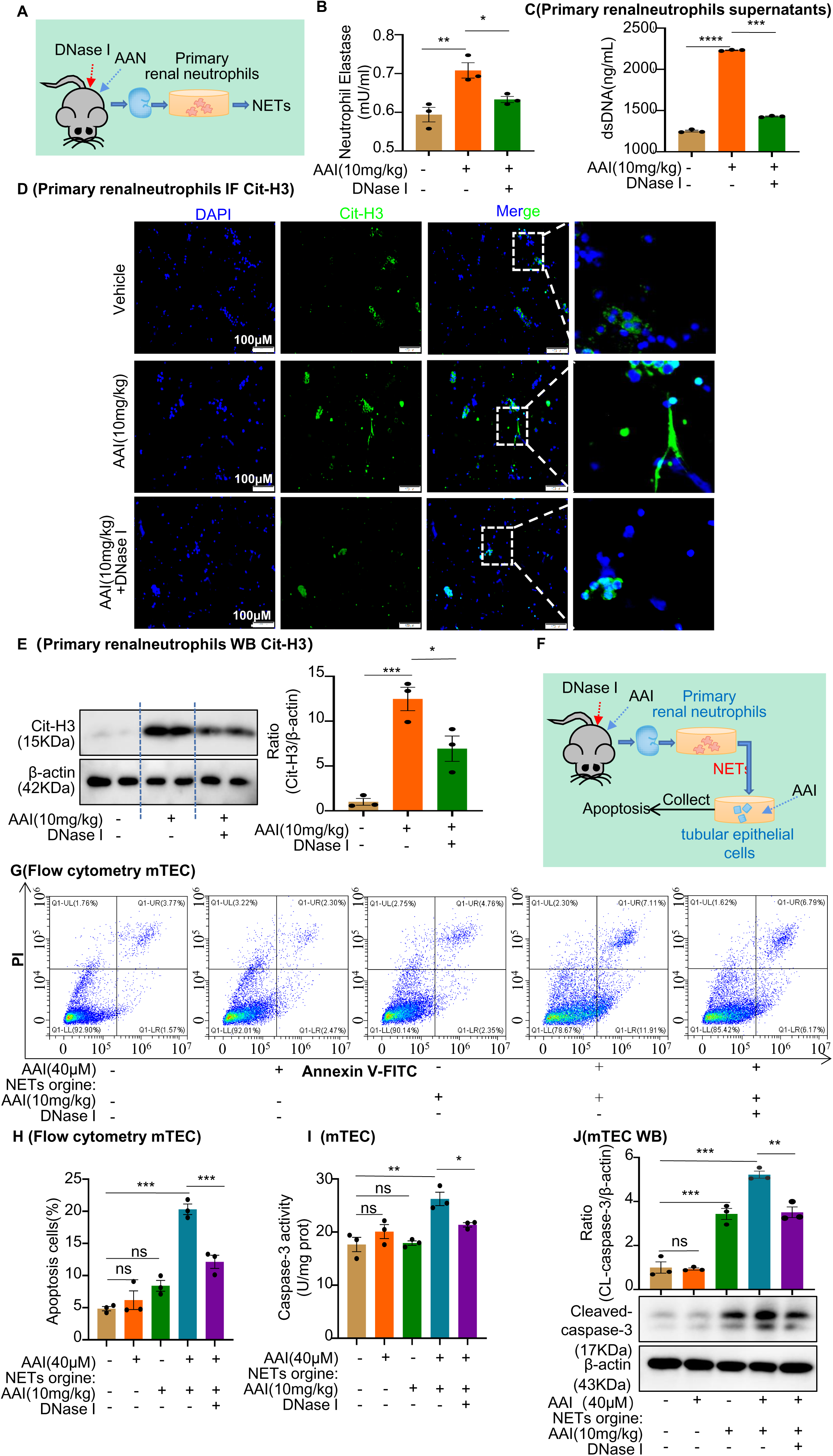
Conditional medium (CM) derived from primary neutrophils of aristolochic acid nephropathy (AAN) mice co-treated with neutrophil extracellular trap (NET) inhibitor prevents apoptosis of mouse renal tubular epithelial cells (mRTECs). **(A)** Primary renal neutrophils were isolated from aristolochic acid I (AAI)- and DNase I-treated mice. **(B)** Level of neutrophil elastase assessed after treating mRTECs with different CMs (n = 3). **(C)** Level of dsDNA assessed after treating mRTECs with different CMs (n = 3). **(D)** Level of citrullinated histone 3 (Cit-H3) assessed using immunofluorescence staining after treatment with different CMs (n = 3). **(E)** Level of Cit-H3 assessed by western blotting after treatment with different CMs (n = 3). **(F)** Primary neutrophil/mRTEC-conditioned media culture model (n = 3). **(G)** Apoptosis in mRTECs cultured with different CMs detected using flow cytometry (n = 3). **(H)** Quantity of apoptotic mRTECs cultured with different CMs (n = 3). **(I)** Activity of caspase-3 quantified after treating mRTECs with different CMs (n = 3). **(J)** Protein levels of cleaved caspase-3 assessed by western blotting after treatment with different CMs (n = 3). Data are presented as the mean ± SEM of three biological replicates per condition. Each dot represents a sample. Statistically significant differences were determined by an independent sample *t*-test and one-way ANOVA followed by Tukey’s post hoc test. *P < 0.05, **P < 0.01, ***P < 0.001, ****P < 0.0001, ns: non-significant. **Figure S8—source data 1** **Figure S8—source data 2** **Conditional medium (CM) derived from primary neutrophils of aristolochic acid nephropathy (AAN) mice co-treated with neutrophil extracellular trap (NET) inhibitor prevents apoptosis of mouse renal tubular epithelial cells (mRTECs).**

**Figure S9.**
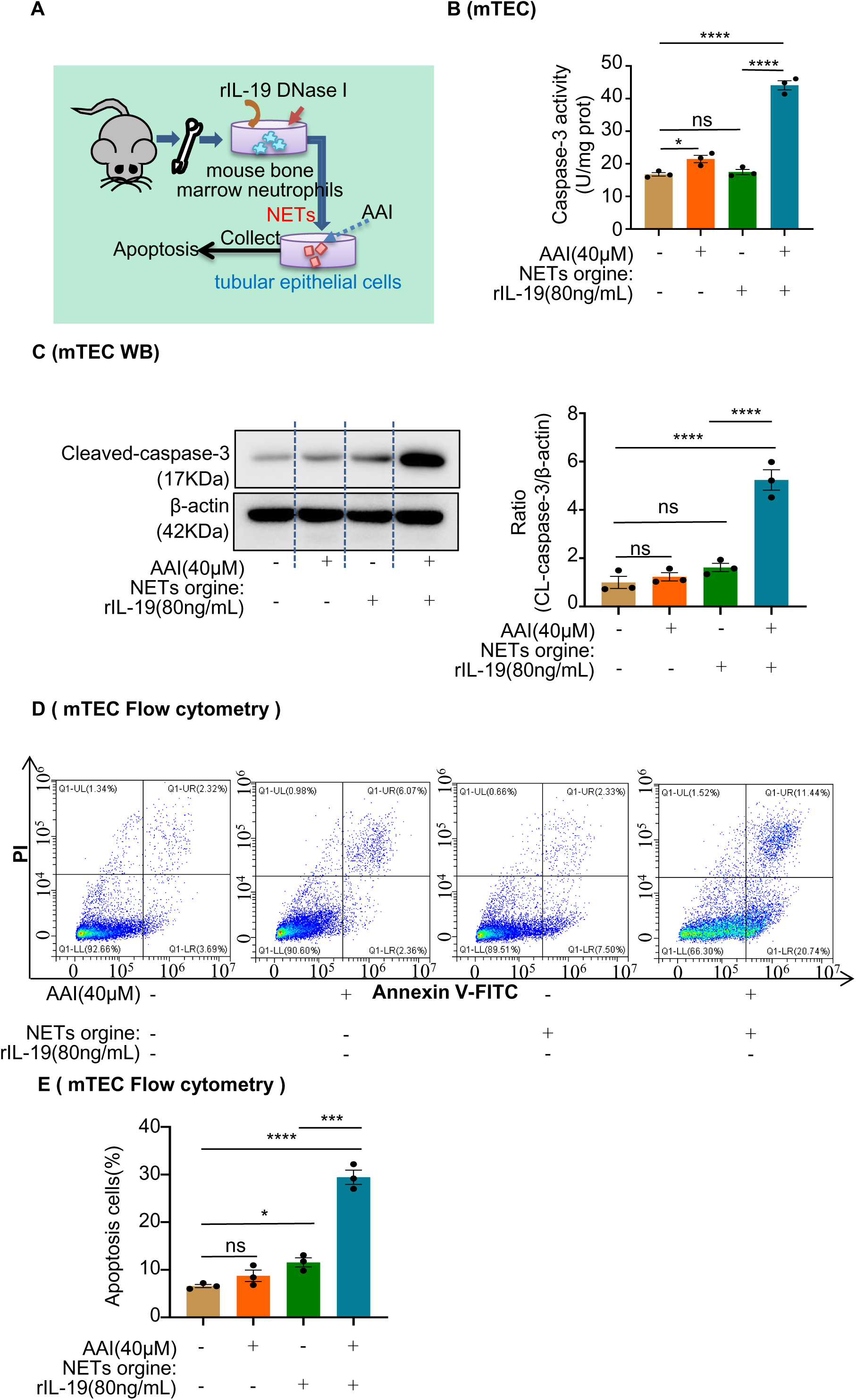
Expression of interleukin-19 (IL-19) in mouse kidneys and mouse tubular epithelial cells (mRTECs) in aristolochic acid I (AAI)-induced aristolochic acid nephropathy (AAN). **(A)** Immunofluorescence (IF) staining of IL-19 (red) and E-cadherin (green) in kidney tissues of AAI-induced AAN mice. E-cadherin was used as an epithelial cell marker (mRTECs, n = 6). Scale bar, 50 μm. **(B)** IF staining of IL-19 (red) in AAI-treated *PSTPIP2*^Flox/Flox^ (FF) and *PSTPIP2*-cKI mice (n = 6). Scale bar, 50 μm. **(C)** IF staining of IL-19 (red) in AAI-treated *PSTPIP2-*cKI mice with AAV-EV or AAV-OE (n = 6). Scale bar, 50 μm. **(D and E)** Verification of *PSTPIP2* overexpression in mRTECs (n = 3). **(F)** IF staining of IL-19 in AAI-treated mRTECs overexpressing *PSTPIP2* (n = 3). **(G)** Bone marrow-derived neutrophils isolated from C57BL/6J mice were identified by flow cytometry. Data are presented as the mean ± SEM of three biological replicates per condition. Each dot represents a sample. Statistically significant differences were determined by an independent sample *t*-test and one-way ANOVA followed by Tukey’s post hoc test. *P < 0.05, ***P < 0.001, ****P < 0.0001, ns: non-significant. **Figure S9—source data 1** **Figure S9—source data 2** **Expression of interleukin-19 (IL-19) in mouse kidneys and mouse tubular epithelial cells (mRTECs) in aristolochic acid I (AAI)-induced aristolochic acid nephropathy (AAN).**

**Figure S10.**
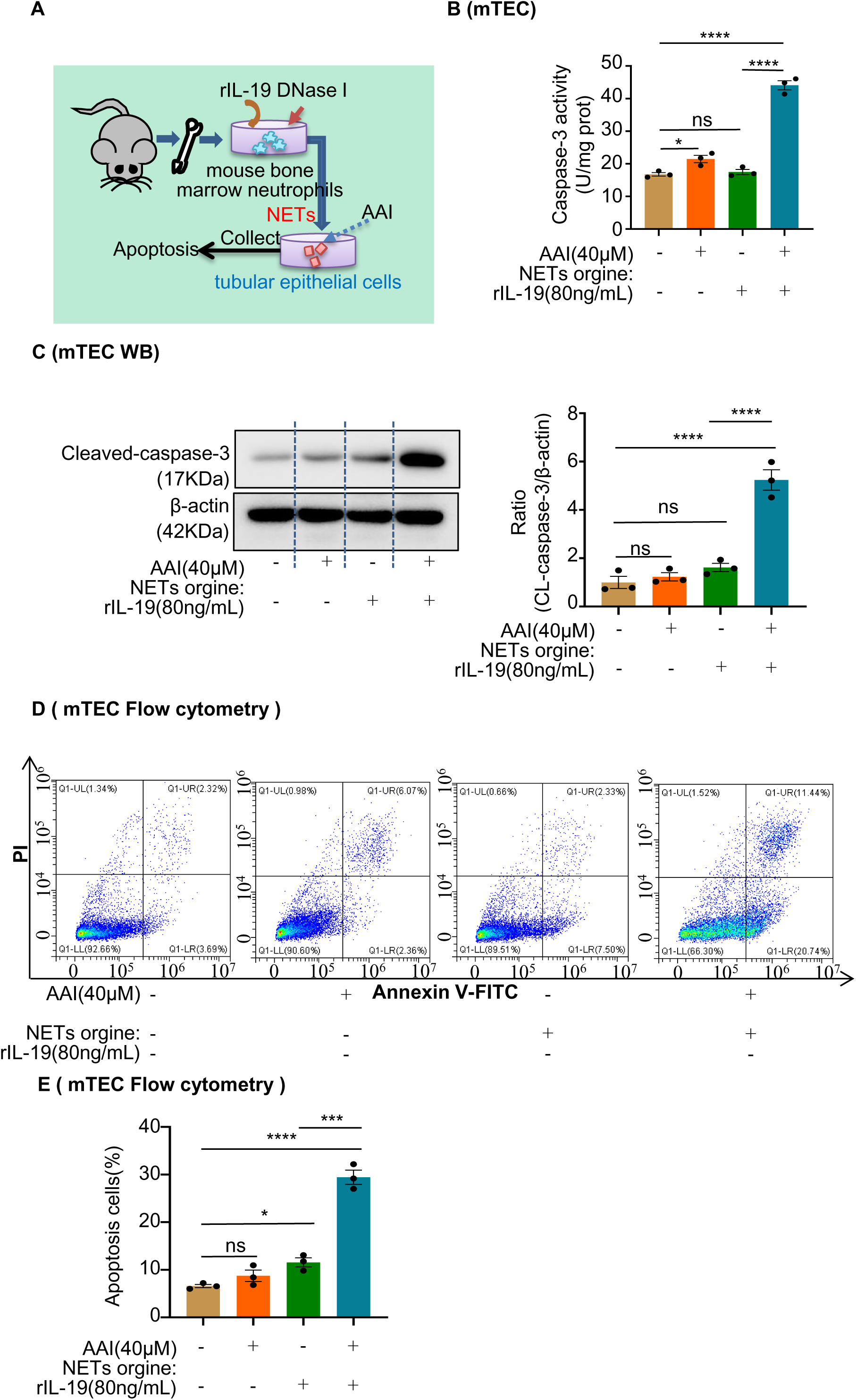
Conditional medium (CM) derived from mouse bone marrow-derived neutrophils co-treated with interleukin-19 (IL-19) promotes apoptosis of injured mouse renal tubular epithelial cells (mRTECs). **(A)** Mouse bone marrow-derived neutrophil/mRTEC-conditioned medium culture model. **(B)** Activity of caspase-3 quantified after treatment with different CMs (n = 3). **(C)** Protein levels of cleaved caspase-3 assessed by western blotting after treatment with different CMs (n = 3). **(D)** Level of apoptosis in mRTECs treated with different CMs detected using flow cytometry (n = 3). **(E)** Quantity of apoptotic mRTECs cultured with different CMs (n = 3). Data are presented as the mean ± SEM of three biological replicates per condition. Each dot represents a sample. Statistically significant differences were determined by an independent sample *t*-test and one-way ANOVA followed by Tukey’s post hoc test. *P < 0.05, ***P < 0.001, ****P < 0.0001, ns: non-significant. **Figure S10—source data 1** **Figure S10—source data 2** **Conditional medium (CM) derived from mouse bone marrow-derived neutrophils co-treated with interleukin-19 (IL-19) promotes apoptosis of injured mouse renal tubular epithelial cells (mRTECs).**

